# *MAPT* Splicing Modulators as a Therapeutic Strategy for Tauopathies

**DOI:** 10.1101/2025.05.06.652428

**Authors:** M. Catarina Silva, Hannah Lindmeier, Paolo Pigini, Jennifer Laughlin, Yong Yu, Christie Morrill, Michael A. Arnold, Scott J. Barraza, Khalil Saadipour, Monal Dieterich, Angela Minnella, Kausiki Datta, Nanjing Zhang, Jana Narasimhan, Christopher R. Trotta, Matthew G. Woll, Ellen M. Welch, Kellie Benzow, Kul Karanjeet, Steven Lotz, Taylor Bertucci, Sally Temple, Stephen J. Haggarty, Michael Koob, Jeffrey Trimmer, Marla Weetall, Elisabetta Morini

## Abstract

Tauopathies are neurodegenerative diseases characterized by the abnormal accumulation of microtubule-associated protein tau (MAPT) in the brain. These disorders, like frontotemporal dementia (FTD-Tau), currently lack effective therapies and can occur sporadically or be inherited when associated with *MAPT* gene mutations. The *MAPT* gene region encompassing exon 10 and adjacent introns is a hotspot for pathogenic variants, including splicing mutations that enhance exon 10 inclusion and increase 4R tau expression, and gain-of-function mutations that generate aggregation-prone mutant 4R tau protein. For these 4R-specific tauopathies, a targeted mRNA splicing approach that promotes exon 10 exclusion may offer therapeutic benefit. In this study, we discovered novel splicing modulator compounds (SMCs) that promote *MAPT* exon 10 exclusion, and demonstrated their efficacy in FTD patient-derived neuronal models carrying the tau-P301L gain-of-function mutation or the tau-S305N splicing mutation. Treatment with SMC reduced 4R tau expression and decreased the accumulation of hyperphosphorylated tau (pTau), oligomeric and insoluble tau, thereby rescuing tau-associated neuronal toxicity. Importantly, our lead SMC corrected the 3R/4R splice ratio *in vivo* and significantly reduced pTau in the brain of a gene-replacement (GR) mouse model expressing the human tau-N279K splicing mutation. These findings support the therapeutic potential of this class of small molecules and establish *MAPT* pre-mRNA splicing modulation as a promising strategy for the treatment of 4R tauopathies.

**One Sentence Summary:** Discovery of SMCs that correct *MAPT* splicing, reduce 4R tau, and rescue pathology in patient-derived neuronal and *in vivo* models of 4R tauopathies.

## INTRODUCTION

Tauopathies are a group of clinically heterogeneous neurodegenerative diseases characterized by the aberrant accumulation of microtubule-associated protein tau (MAPT) in the form of oligomers, aggregates, neurofibrillary tangles, and paired helical filaments in neurons and glia of the affected brain regions. Disorders like frontotemporal dementia with tau pathology (FTD-Tau, hereafter referred to as FTD) lack effective therapies and may manifest in either sporadic or inherited forms, the latter linked to mutations in the *MAPT* gene at chromosome 17q21.3 (*1, 2*). In the human brain, six tau isoforms are expressed through a developmentally regulated process involving alternative splicing of *MAPT* exons 2, 3, and 10 (Fig. 1A) (*3, 4*). Alternative splicing of exons 2 and 3 produces the 0N, 1N, and 2N tau isoforms of the N-terminal projection region (Fig. 1A), which play a role in signal transduction and membrane interactions (*3*). Exon 10 encodes the second microtubule (MT)-binding repeat domain (R2) in the C-terminal region (Fig. 1A). Inclusion of exon 10 leads to expression of 4R tau, with four MT-binding repeat domains, whereas exclusion of exon 10 leads to expression of 3R tau (*4, 5*). The MT-binding repeat domains (Fig. 1A) are essential for tau to regulate the stability and dynamics of microtubules as well as support axonal transport, with 4R tau isoforms binding to MTs with greater affinity than 3R tau isoforms; for this reason, the relative 4R/3R expression is developmentally regulated (*3, 4*). The main isoform present in the fetal stage is 3R tau, when the dynamic nature of axons and neuronal projections is essential for synaptogenesis and the development of neural networks. In the adult healthy brain, the overall 4R/3R ratio is maintained at 1 (Fig. 1B) (*4*).

**Figure 1.**
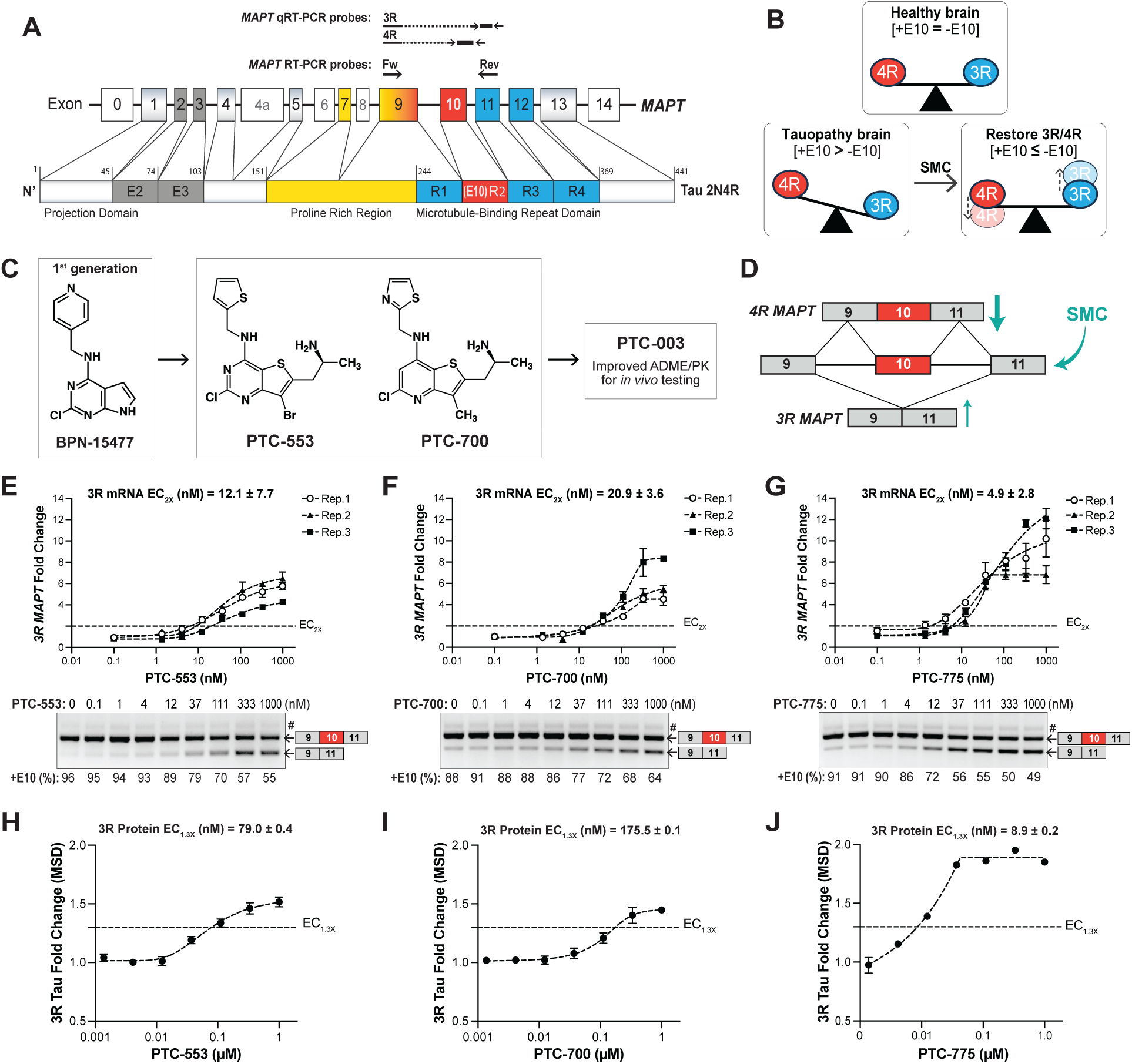
Identification of splicing modulator compounds that promote *MAPT* exon 10 exclusion. **(A)** Schematic of the human *MAPT* gene exons and longest tau protein isoform (2N4R). Exon 10 and the encoded MT-binding repeat domain R2, which can be alternatively spliced to generate 3R (–E10) and 4R (+E10) tau isoforms, are highlighted in red. Primers binding sites used for RT-(q)PCR are indicated above the schematic. **(B)** Schematic representation of 4R/3R tau ratio in health and disease. In the human adult brain, 4R/3R tau ratio is 1, with both isoforms found at similar levels; however, in specific forms of tauopathy (*bottom left*) there is an increase in 4R tau expression (+E10) via exon 10 splicing inclusion and/or a propensity for 4R tau accumulation. Our goal is to identify SMCs that rescue the balance of 4R/3R ≃ 1 or even reduce the expression of 4R so that 4R/3R < 1 (*bottom right*). **(C)** SAR-based optimization of BPN-15477 to generate more potent splicing modulators. Compounds were screened based on their ability to increase *MAPT* exon 10 skipping, assessed by qRT-PCR and protein analysis. **(D)** Schematic of the reporter minigene assay to assess *MAPT* exon 10 splicing. Numbered boxes represent exons 9, 10 (red), and 11. The *4R* transcript (*top*) results from exon 10 inclusion, while the *3R* transcript (*bottom*) results from exon 10 skipping, a process enhanced by a splicing modulator compound (SMC). **(E-G)** Minigene assay in HEK293T cells stably expressing the human *MAPT* gene, treated with PTC-553 (E), PTC-700 (F) and PTC- 775 (G) for 24h. RT-qPCR analysis was performed to measure exon 10 skipping and the resulting increase in *3R MAPT* levels. EC_2X_ represents the compound effective concentration inducing a 2-fold increase in *3R MAPT* relative to baseline. Data points represent *3R* fold-change relative to DMSO-treated cells ± SEM, with *N* = 3 biological replicates. Representative RT-PCR gel electrophoresis of *MAPT* splicing below each graph shows dose-dependent increase in *3R MAPT* (–E10, lower band) and a corresponding decrease in *4R MAPT* (+E10, upper band). A faint higher band (#) shares the same nucleotide sequence as the *4R MAPT* band, likely representing a secondary structure of the PCR product. **(H-J)** MSD assay to measure changes to 3R tau protein in U-87 glioma cells. Data points represent mean fold-change relative to DMSO-treated cells ± SEM with *N* ≥ 3. EC_1.3X_ is annotated and refers to the effective concentration of each compound inducing a 1.3-fold increase in 3R tau relative to DMSO treatment.

Neuropathological evidence shows that tau inclusions from different tauopathies have different tau isoform compositions, suggesting a key role for splicing and isoform-specific contribution to pathophysiology (*5, 6*). Whereas *MAPT* mutations affecting exons 2 and 3 are rare, exon 10 and its intronic boundary regions are a hot spot for pathogenic variants, representing approximately 27% of all known tau mutations (*1, 5, 6*). The majority of mutations in intron 10 are clustered in the stem-loop region (*7, 8*). These can alter the RNA structure and accessibility of the spliceosome, resulting in mis-splicing, which in most cases increases inclusion of exon 10 and enhances 4R tau expression, which has a higher propensity for aggregation (*9, 10*). Splicing mutations that cause an imbalance in 4R/3R tau expression have been associated with several forms of FTD (*2, 9–13*) (Fig. 1B). Prominent examples include the autosomal dominant S305N missense mutation in the last codon of *MAPT* exon 10, within the RNA hairpin-loop structure (*12, 14, 15*), and the autosomal dominant N279K missense mutation also in exon 10, identified as one of the most frequent causes of familial FTD (*11, 13*). In a different category, autosomal dominant gain-of-function mutations in exon 10, like tau-P301L, do not affect splicing or isoform expression ratio but result in the production of mutant 4R tau protein with an elevated propensity for aberrant post-translational modifications, misfolding, and accumulation in the form of aggregates and filaments (*16–18*). Tau-P301L is one of the most common mutations associated with 4R tauopathies worldwide (*9, 16, 18*). In both contexts, a therapeutic approach that promotes *MAPT* exon 10 exclusion might be beneficial (Fig. 1B).

To date, there are no approved disease-modifying therapies for tauopathies, and few tau-targeting drugs have reached clinical trials (*19–28*). Current experimental therapeutics include small-molecule aggregation inhibitors, inducers of tau clearance, and antibody-based immunotherapies targeting aggregated, fibrillary, or seed-competent tau (*22–24, 29–36*). Additionally, RNA-based modalities such as small interfering RNA (siRNA) and antisense oligonucleotides (ASOs) can reduce total or 4R *MAPT* expression, and decrease tau accumulation, prevent neuronal loss, and rescue behavioral phenotypes in mouse and non-human primate models (*20, 21, 37–39*), supporting *MAPT* splicing as a relevant disease-modifying target. However, ASOs for tau have faced some barriers, including low brain and cellular permeability, toxicity associated with immunogenicity, and invasive routes of administration (*40–43*). Small-molecule splicing modulator compounds (SMCs) are a promising alternative, supported by the recent advances in the development of SMCs for neurological disorders such as spinal muscular atrophy (SMA) and Huntington’s disease (*44–49*). For tau, innovative strategies include small molecules directly targeting unique structural features of the *MAPT* pre-mRNA that have been shown to lower the 4R/3R tau ratio *in vivo* (*14, 50*). *MAPT* splicing modulators offer several advantages by directly targeting the underlying cause of the disease, enabling post-transcriptional intervention without genome editing, and allowing dose-dependent target engagement. SMCs also have additional benefits, including favorable systemic and brain distribution, easy delivery route, the possibility of oral administration, and rapid target engagement with splicing changes occurring within hours of administration (*51*).

Our team initially developed SMCs for familial dysautonomia (FD), a rare neurodegenerative disease caused by a splicing mutation in the *ELP1* gene (*52–56*). One such compound, BPN-15477 (Fig. 1C), was identified through a machine learning-based approach to also promote *MAPT* exon 10 exclusion (*57*). Notably, BPN-15477 demonstrated high specificity, with transcriptome analysis of treated fibroblast cells revealing changes in exon splicing for only 0.58% of all expressed exon triplets (934 out of 161,097 expressed triplets) (*57*). Given the need for therapies that modulate the 4R/3R tau ratio, we used BPN-15477 as a starting point to develop novel *MAPT* splicing modulators and to evaluate their therapeutic potential in neuronal and mouse models of FTD (*57*).

Patient induced pluripotent stem cell (iPSC)-derived, cortical-enriched neurons offer a unique advantage to studying disease-associated mis-splicing events in a relevant physiological, cellular, and genomic background (*58, 59*). We employed tau-P301L (*36*) and tau-S305N (*15, 60*) FTD patient iPSC-derived neuronal cell models to evaluate the effect of SMC treatment on endogenous *MAPT* splicing, tau protein expression and accumulation, and mutation-associated phenotypes. Furthermore, we tested our lead SMC *in vivo*, in a new transgenic gene-replacement (GR)-mouse model of FTD carrying the human tau-N279K splicing mutation (*MAPT*- GR*N279K) (*61*). Our findings demonstrate that SMC treatment can significantly reduce 4R tau expression in FTD patient iPSC-derived neurons and *in vivo*, thereby reducing tau accumulation and toxicity. Overall, these results highlight the druggability of splicing regulation and the therapeutic potential of these compounds as a novel strategy for treating 4R tauopathies.

## RESULTS

### Development of highly potent *MAPT* splicing modulator compounds

We identified BPN-15477 (Fig. 1C) as a splicing modulator of *MAPT* exon 10, using a machine learning-based approach to identify gene targets for this novel class of SMCs (*62*). BPN-15477 promoted *MAPT* exon 10 exclusion in HEK293T cells stably expressing a *MAPT* minigene (*62*) (Fig. 1D). To validate the effect of BPN-15477 on endogenous human *MAPT* splicing and in a neuronal context, we tested this compound in FTD patient iPSC-derived neurons expressing the 4R-specific gain-of-function mutation tau-P301L (*36*). Neurons were treated with compound concentrations between 10 μM and 100 μM during the final 5 weeks of an 8-week differentiation period (Fig. S1). BPN-15477 reduced 4R tau expression at the transcript (Fig. S1A) and protein levels (Fig. S1B) but also caused some toxicity, as shown by the dose-dependent reduction in the levels of neuronal markers β-III-tubulin (axonal protein, TUJ1) and PSD95 (synaptic protein) (Fig. S1C). This was not surprising given the high concentrations of BPN-15477 (50–100 μM) required to observe an effect on *MAPT* splicing, likely due to its low potency.

To develop more potent SMCs capable of effectively modulating *MAPT* splicing in the brain, we established a structure-activity relationship (SAR) by generating analogs of BPN-15477. We discovered structural modifications that afforded analogs with improved potency, efficacy, and drug-like properties (Fig. 1C). To evaluate compounds that modulate *MAPT* exon 10 splicing, we used a HEK293T cell line stably expressing a *MAPT* minigene that comprises exon 10, the 3’- end of exon 9, the 5’-end of exon 11, and ∼150 nucleotides of the flanking introns of the human gene (*62*) (Fig. 1D). This system was specifically optimized for compound profiling, as it overcomes the challenge of low endogenous expression of *4R MAPT* transcript in most cell lines. The minigene is predominantly spliced to include exon 10, resulting in a high baseline level of *4R MAPT* expression. Treatment with SMCs that promote exon 10 skipping leads to increased production of the *3R MAPT* transcript. In this study, we present findings on three lead compounds: PTC-553 (Fig. 1C, E), PTC-700 (Fig. 1C, F), and PTC-775 (Fig. 1G), which promote a dose- dependent increase in *3R MAPT* as measured by RT-qPCR and complementary RT-PCR gel analysis (Fig. 1E-G). For these compounds, the effective concentration inducing a 2-fold change (EC2X) relative to vehicle-treated neurons was between 5 nM and 21 nM after 24h of treatment. For subsequent validation, we utilized a Meso Scale Discovery (MSD)-based assay to measure compound-induced changes in endogenous 3R tau protein in a human U-87 glioma cell line. This plate-based technology provides ultra-sensitive measures similar to traditional ELISA but with improved sensitivity. MSD results showed a compound dose-dependent increase in endogenous 3R tau protein (Fig. 1H-J), reflecting the increase in *MAPT* exon 10 skipping. In this case, the calculated EC1.3X, which represents the threshold concentration required to achieve a near-maximal effect (lower effective concentration indicates greater potency), was also in the nM range. Based on these results, we ranked our SMCs by EC2X for *3R* mRNA and EC1.3X for 3R protein, with PTC- 775 exhibiting the highest potency *in vitro*, followed by PTC-553 and PTC-700 (Fig. 1E-J).

### 4R tauopathy modeled in FTD patient iPSC-derived neurons

To study the therapeutic effect of SMCs for tauopathies, we employed two FTD patient iPSC-derived neuronal models. The first model is derived from a carrier of the autosomal dominant, gain-of-function tau-P301L missense mutation (*35, 36*), which leads to the expression of 4R- specific mutant tau (Fig. 2A). The second model originates from a patient carrying the autosomal dominant S305N missense splicing mutation (*15, 60*), which increases the expression of *4R MAPT* (Fig. 2A). Whereas we have comprehensively characterized the previously published P301L neuronal model (*35, 36, 63*), the tau-S305N iPSC-derived neuronal model was newly generated and characterized following published methodologies (*15, 60*). Briefly, within 8 weeks of neural progenitor cells (NPCs) differentiation (Fig. S2A), we observed elevated *4R MAPT* mRNA levels in S305N neurons by RT-PCR, relative to *3R MAPT*, and in contrast to very low *4R MAPT* levels in control (wild type/WT tau) and P301L neurons (Fig. S2B). We then evaluated the differentiation profile of tau protein in S305N neurons, in parallel with WT and P301L neurons, during 8 weeks from the NPC stage. As expected, tau levels increased in a differentiation-dependent manner, with notable differences among the three genotypes (Fig. S2C-E). As previously described (*35, 36*), 6- and 8-week P301L neurons exhibit a more rapid increase in total tau (TAU5 antibody, Fig. S2D) and phosphorylated tau (pS396, pT231, and pS202/T205), including an increase in detection of high molecular weight (MW) oligomeric pTau-S396, compared to non-mutant, healthy control neurons (Fig. S2C, E). This reflects the accumulation of tau species prone to misfolding and oligomerization. However, 3R and 4R tau levels remained similar between P301L and control neurons during this differentiation period (Fig. S2D). In turn, S305N neurons showed a more rapid increase in 4R tau (Fig. S2C, D) consistent with mutation-driven enhanced exon 10 inclusion and increased *4R MAPT* transcript expression (Fig. S2B). The faster accumulation of pTau was also significant and included the accumulation of high MW oligomeric pS396 tau species of reduced SDS solubility after 6 weeks of differentiation (Fig. S2C, E). As an additional control for this study, total tau (TAU5) and isoform-specific tau antibodies were validated in protein lysates from human adult brain tissue and purified recombinant tau isoforms (Fig. S2F), reinforcing confidence in the detection of 4R tau in iPSC-derived neuronal lysates. The distinct tau profiles observed between P301L and S305N neuronal differentiation (Fig. S2D, E) underscore the unique molecular impact of these mutations, supporting their selection for evaluating the effects of SMCs in FTD.

**Figure 2.**
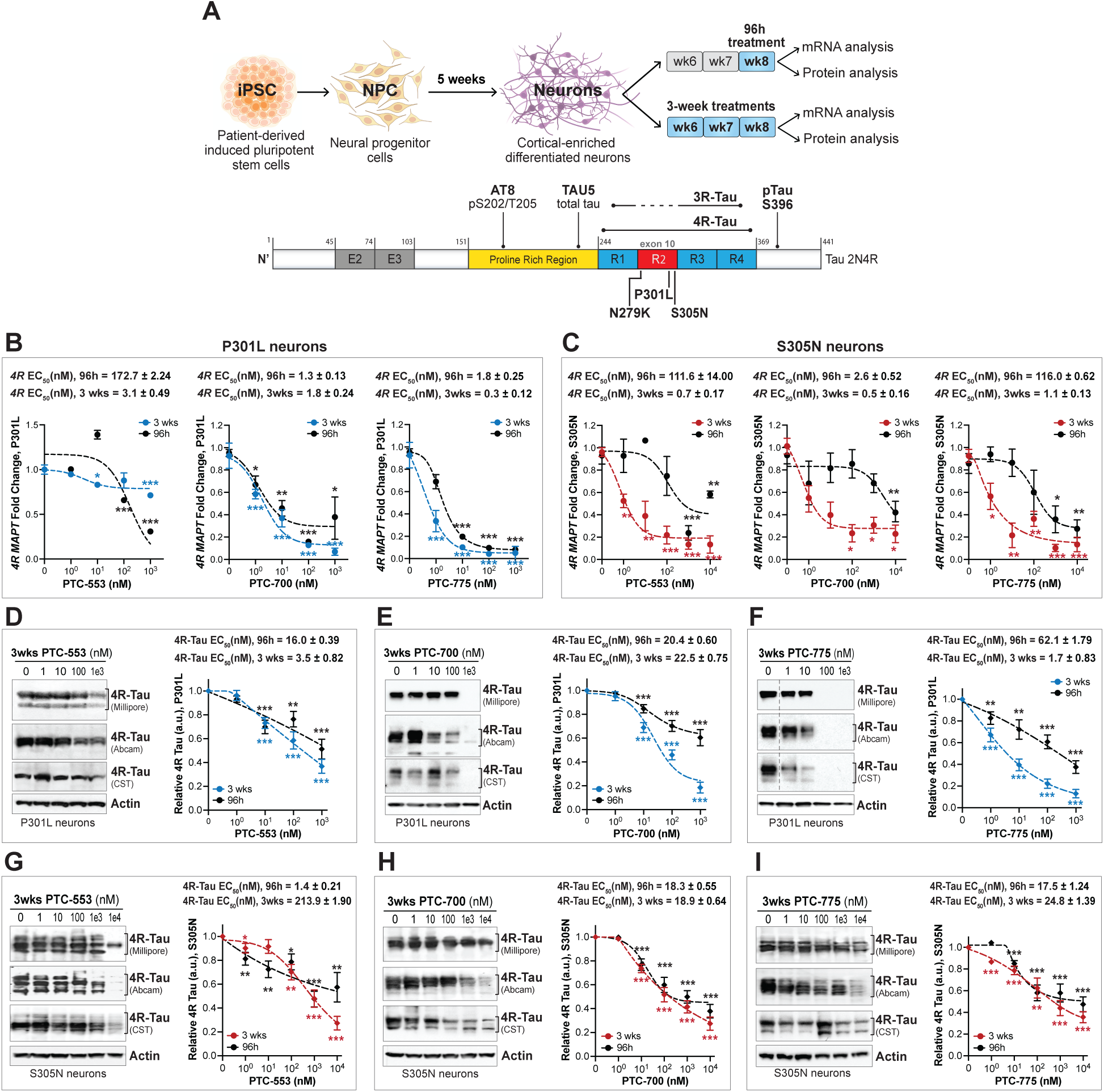
SMCs reduce *4R MAPT* expression and 4R tau protein levels in patient iPSC-derived tauopathy neuronal models. **(A)** Schematic representation of iPSC-to-NPC and neuronal differentiation and compound treatment regimens: single 96h treatment during week 8 of differentiation, or weekly treatment from weeks 5 to 8, i.e., 3-week treatment. Below, the 2N4R tau isoform is shown, highlighting exon 10 (MT-binding domain R2), a hotspot for pathogenic variants associated with tauopathy. Also indicated are the main total tau and pTau antibodies used in this study. **(B)** RT-qPCR analysis of SMCs effect on *4R MAPT* in tau-P301L neurons and in **(C)** tau-S305N neurons following a 96h or 3-week treatment. EC_50_ for *4R MAPT* are indicated. Data points represent mean values relative to DMSO-treated neurons (normalized to *GAPDH*) ± SEM, with N ≥ 3 biological replicates. **(D-F)** Western blot analysis of SMCs dose-effect on 4R tau after 96h or 3 weeks of treatment of P301L neurons, with representative blots of the 3-week treatment shown for each compound. Densitometry analysis of 4R tau protein, relative to DMSO-treated neurons (normalized to actin) is shown. **(G-I)** Corresponding analysis for S305N neurons. Blot images show results obtained with three different 4R-specific antibodies, while the graphs show the average quantification for the three antibodies. Dashed lines indicate cropped images solely for the purpose of this figure to exclude data unrelated to this study, all samples were run in the same gel. EC_50_ for 4R tau are indicated. Data points represent mean densitometry relative to DMSO-treated neurons ± SEM, with N ≥ 3. Statistical analysis (B-I) was performed using a two-tailed, paired Student’s t-test for each compound concentration vs. vehicle (0 μM). Significance thresholds: \**p* ≤ 0.05, \*\**p* ≤ 0.01, and \*\*\**p* ≤ 0.001.

### SMCs significantly reduce 4R tau in P301L and S305N FTD patient iPSC-derived neurons

To assess the effect of SMCs on human neuronal *MAPT* transcript expression and tau protein, we employed 8-week differentiated P301L and S305N neurons (Fig. 2A). This choice was based on the observed increase in tau and pTau accumulation between weeks 6 and 8 of differentiation, as well as the upregulation of 4R tau in S305N neurons during the same period (Fig. S2B-E), both representing 4R tau-dependent phenotypes. To account for the active differentiation and maturation of the neuronal cultures at this stage, and the continued increase in 4R tau expression, we implemented two treatment regimens (Fig. 2A): a 96h treatment during week 8 of differentiation, and a weekly treatment from weeks 5 to 8 of differentiation, representing a longer, 3-week multi-treatment.

To evaluate the effect of the lead SMCs PTC-553, PTC-700, and PTC-775 on exon 10 skipping, we measured *4R MAPT* mRNA levels by RT-qPCR in P301L (Fig. 2B) and S305N (Fig. 2C) neurons treated for 96h or 3 weeks. Since 8-week-old iPSC-derived neurons express high levels of *3R MAPT* (particularly in P301L neurons), monitoring exon 10 skipping via changes in *4R MAPT* provided greater sensitivity to small-level changes. In both neuronal models, we observed a dose-dependent decrease in *4R MAPT* upon SMC treatment (Fig. 2B, C). Additionally, we noted a general trend toward higher SMC potency with the 3-week treatment, accompanied by a decrease in the respective half-maximal effective concentration, or EC50, for *4R MAPT* reduction. Based on EC50 values, which reflect compound potency, PTC-553 was the weaker compound, showing no clear dose-dependent effect on *4R MAPT* in P301L neurons (Fig. 2B), and only exhibiting a dose-dependent response in S305N neurons after the 3-week treatment (EC50 = 0.15 nM, Fig. 2C). PTC-700 demonstrated greater potency, with a clear dose-dependent reduction in *4R MAPT* in both P301L (EC50 = 1.3-1.8 nM, Fig. 2B) and S305N neurons (EC50 = 0.5-2.6 nM, Fig. 2C). PTC-775, an improved analog of PTC-700, induced the strongest reduction in *4R MAPT* at the highest concentrations, with an EC50 of 0.3-1.8 nM in P301L neurons (Fig. 2B) and 1.1-116.0 nM in S305N neurons (Fig. 2C). In S305N neurons, because this mutation significantly increases expression of *4R MAPT* in iPSC-derived neurons (Fig. S2B), we were able to also show by RT-PCR gel electrophoresis analysis of *MAPT* splicing, a dose-dependent increase in *3R MAPT* (–E10, lower band) and a corresponding decrease in *4R MAPT* (+E10, upper band) (Fig. S3A-C), which corroborated the initial minigene screening data (Fig. 1E-G). In summary, all SMCs effectively reduce *4R MAPT* expression in human neurons at nanomolar concentrations, with the extent of *4R* reduction correlating with SMC potency.

Next, we sought to determine the extent of SMC effect on 4R tau protein in the FTD patient iPSC-derived neurons. To this end, we tested and optimized detection of 4R tau using three commercially available antibodies (see Fig. S3D-F for P301L and Fig. S3G-I for S305N individual antibody results). In P301L neurons, the three SMCs induced a dose-dependent reduction in 4R tau after 96h of treatment, with EC50 values ≤ 62 nM (Fig. 2D-F). We also observed a trend for improvement in 4R tau reduction and lower EC50 values upon 3 weeks of treatment (Fig. 2D-F: blue vs. black curves). PTC-775 showed the highest potency (up to 90% reduction) and strongest dose-dependent 4R tau reduction, particularly upon 3 weeks of treatment (EC50 = 1.7 nM, Fig. 2F). It is followed by PTC-700 (EC50 = 20.4-22.5 nM, Fig. 2E) and PTC-553, which presents with low EC50 (3.5-16 nM), but achieved the weakest reduction in 4R (∼ 50%, Fig. 2D). The results in S305N neurons were equally positive (Fig. 2G-I). PTC-700 (Fig. 2H) and PTC-775 (Fig. 2I) showed the highest potency (EC50 = 17-25 nM), and an improved effect upon longer treatment, leading to a strong reduction in 4R tau by 60-80% relative to vehicle-treated neurons (red curves, Fig. 2H, I). PTC-553 exhibited a dose-dependent effect with lower EC50 at 96h (1.4 nM, Fig. 2G), but improved effect after 3 weeks of treatment, lowering 4R tau by 80% (red curve, Fig. 2G).

Overall, the SMCs effect on 4R tau protein was closely related to the effects observed at the transcript level, replicating the compound ranking with PTC-700 and PTC-775 emerging as top leads. These findings highlight the efficacy of our SMCs in reducing 4R tau at both the transcript and protein levels, indicating potential to alleviate 4R-associated pathology.

### SMCs reduce hyperphosphorylated tau in FTD iPSC-derived neurons without compromising cell viability

Having demonstrated 4R tau reduction with nanomolar potency in patient-derived neurons, we next assessed SMC effects on neuronal viability. After 96h of treatment, no significant impact on neuronal viability was observed within the effective concentration range tested, for either P301L or S305N neurons (Fig. 3A-C). In 3-week treated P301L neurons (Fig. S4A-C, *black curves*), no significant changes in cell viability were observed, except for a slight trend toward reduced viability at the highest concentration (10 μM), well above the nanomolar effective range. To avoid potential artifacts in data interpretation (*i.e.,* tau reduction as a consequence of neuronal loss), from here on we limited treatments in P301L neurons to a maximum 1 μM concentration, as this was sufficient to achieve a significant reduction in 4R tau at both the transcript and protein levels (Fig. 2). Interestingly, S305N neurons treated for 3 weeks (compound added to cell medium once per week), showed improved cell viability (10-20% increase) with 1–10 μM SMC concentrations (Fig. S4A-C, *blue curves*), further underscoring the value of using these two tau models to study the effects of SMC treatment.

**Figure 3.**
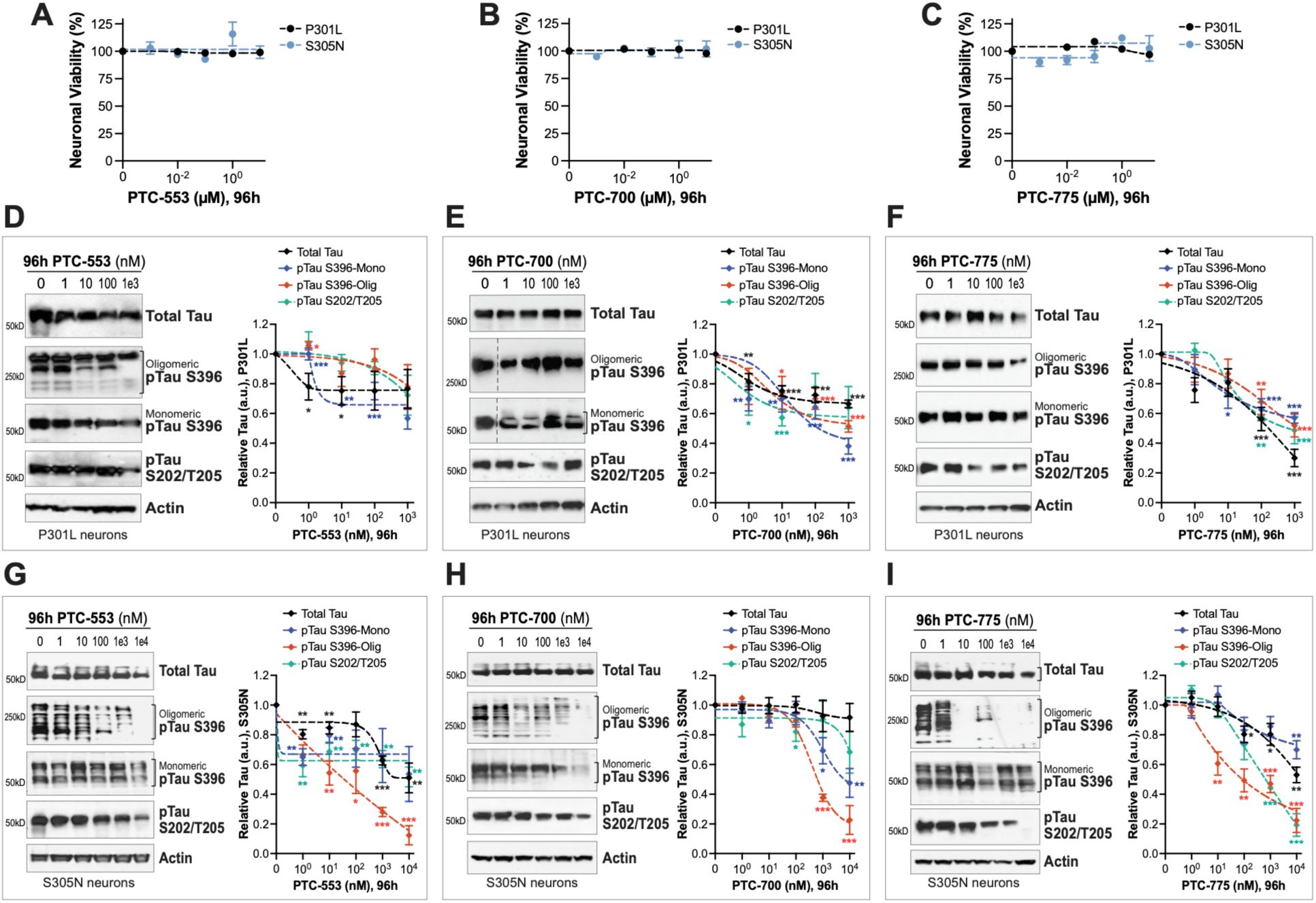
SMC treatment reduces accumulation of total tau and pTau without compromising neuronal viability. (A-C) Viability dose curves for 8-week differentiated P301L and S305N neurons treated with PTC-553 (A), PTC-700 (B) or PTC-775 (C) for 96h. Data points represent mean viability relative to DMSO-treated neurons (100% viability) ± SEM, with *N* = 3 biological replicates. **(D-F)** Compounds dose curves for total tau (TAU5 antibody), pTau S396 (high MW oligomeric and monomeric pTau), and pTau S202/T205 (AT8 antibody) after 96h of treatment of P301L neurons, measured by western blot and densitometry analysis. Dashed lines on blots (E) indicate cropped images solely for the purpose of this figure to exclude data points unrelated to this study, all samples were run in the same gel. **(G-I)** Corresponding analysis for S305N neurons treated for 96h. Representative blots are shown. Data points represent mean densitometry relative to DMSO-treated neurons ± SEM, with *N* = 3. Statistical analysis was performed using a two-tailed, paired Student’s t-test for each compound concentration vs. vehicle (0 μM). Significance thresholds: \**p* ≤ 0.05, \*\**p* ≤ 0.01, and \*\*\**p* ≤ 0.001.

To ensure a rigorous assessment of other potential toxic effects by SMC treatment, we measure the levels of key neuronal markers such as NeuN (neuronal nuclear-specific protein), β- III-tubulin (microtubule-associated protein, TUJ1), SYN1 (synapsin 1, synaptic vesicle protein), and PSD95 (postsynaptic density protein 95) using P301L neurons as a test case (Fig. S5). Unlike the first-generation SMC (BPN-15477) that required high μM doses, our SMC lead PTC-700 caused no reduction in the neuronal markers after 96h (Fig. S5A, B) or 3 weeks (Fig. S5F, G) of treatment, indicating no significant toxicity or morphological stress. Similar results were obtained for P301L neurons treated for 96h (Fig. S5K, L) or 3 weeks (Fig. S5P, Q) with PTC-775, although some variability was detected at the highest treatment concentrations that did not reach statistical significance. This is also illustrated by microscopy immuno-cytochemistry (ICC) imaging of P301L neuronal cultures. Images show a clear reduction in pTau-S396/S404 staining (PHF1 antibody; Fig. S5U–pTau in red, see zoom insets) in SMC-treated neurons compared to DMSO-treated controls, with no detectable changes in overall culture density or MAP2-positive dendritic staining patterns (Fig. S5U). Overall, these results confirm that SMCs were well tolerated under the treatment conditions used in this study.

To test the effect of SMCs on tauopathy disease-relevant phenotypes, we started by measuring changes to total tau and pTau protein accumulation after 96h (Fig. 3D-I) or 3 weeks (Fig. S4D-I) of treatment. We focused on pTau-S396, which enables the measurement of monomeric and oligomeric pTau species that are partially SDS-insoluble and migrate as high MW bands in SDS-PAGE (*35, 36, 64*), and pTau-S202/T205 (detected by the AT8 antibody), one of the most widely used markers in brain pathology for tauopathy diagnosis (Fig. 2A). After a 96h treatment with PTC-700 and PTC-775, P301L neurons exhibited a clear dose-dependent reduction in total tau (TAU5), pTau-S396, and pTau-S202/T205 (AT8), achieving a 40–60% reduction of tau at the maximum concentration (Fig. 3E, F). In contrast, PTC-553 showed a limited dose-dependent effect, with a maximum tau reduction of only 20–30% (Fig. 3D). After 3 weeks of treatment, all P301L dose-response curves became more pronounced. PTC-700 (Fig. S4E) and PTC-775 (Fig. S4F) led to 90–100% reduction in total tau and pTau at the maximum concentration. With longer treatment, the effect of PTC-553 also improved, leading to ≥80% reduction in pTau- S396 and ∼40% reduction in total tau and pTau-S202/T205 (AT8) (Fig. S4D). Notably, all SMCs reduced oligomeric pTau-S396 by nearly 100% at the 1 μM concentration, which is expected to have a significant impact on overall tau accumulation and aggregation.

The results in treated S305N neurons were similar. After 96h of treatment, PTC-553 exhibited a dose-dependent reduction for high MW pTau-S396 (Fig. 3G), achieving a 90% reduction at the highest dose of 10 μM, while other markers showed a maximum reduction of 40– 50% with a weak dose-dependent effect. PTC-700 promoted dose-dependent reduction in tau between 0.1 and 10 μM (Fig. 3H), with the strongest reduction (80%) observed for oligomeric pTau-S396. PTC-775 demonstrated the most well-defined dose-response curves (Fig. 3I), achieving an 80% maximum reduction in oligomeric pTau-S396 and pTau-S202/T205 (AT8), potentially the most relevant tau species from a pathological perspective. After 3 weeks of treatment, all SMCs exhibited increased potency. PTC-553 reduced total tau, pTau-S396, and pTau-S202/T205 (AT8) in a dose-dependent manner (Fig. S4G), with the strongest effect seen for oligomeric pTau-S396, which was nearly eliminated at 10 μM (∼100% reduction). PTC-700 showed a similar dose-dependent decrease across all tau markers, with pTau-S202/T205 (AT8) displaying the weakest reduction (∼40%), while total tau and pTau-S396 decreased by up to 80% (Fig. S4H). Finally, PTC-775 showed the strongest reduction for oligomeric pTau-S396 (> 80%) starting at 100 nM, but weaker reduction for pTau-S202/T205 (AT8) and monomeric pS396 (40%) at the maximum dose (Fig. S4I).

Overall, we predict that the observed reduction in total tau steady-state levels after 96h and 3 weeks of treatment in both P301L and S305N neurons results from SMC-mediated mechanisms: decreased expression of mutant 4R tau (P301L) or correction of 4R/3R splicing imbalance (S305N). These effects likely prevent the long-term accumulation of misfolded tau or excess 4R tau, respectively, *i.e.*, the species known to drive tau oligomerization and eventually aggregation. Nonetheless, to rigorously demonstrate that the reduction in tau protein levels was not due to masked toxicity or neuronal loss, we re-analyzed the dose-response curves for 4R tau, total tau, and pTau-S202/T205 (AT8) by normalizing to neuronal markers NeuN and β-III-tubulin (TUJ1), and compared these results to actin-normalized curves in P301L neurons (Fig. S5). NeuN is specifically expressed in neuronal nuclei and β-III-tubulin is a microtubule-specific protein, both providing greater sensitivity for detection of neuronal loss compared to actin alone. For NeuN-normalized data, we observed dose-response profiles for PTC-700 (Fig. S5C, H) and PTC-775 (Fig. S5M, R) treated neurons that were highly similar to the results with actin (Fig. S5E, J, O, T, also see Fig. 3, S4). Normalization to β-III-tubulin yielded comparable results for PTC-700 (Fig. S5D, I vs. E, J) and PTC-775 (Fig. S5N, S vs. O, T). These findings demonstrate that SMC treatment effectively reduces tau accumulation while maintaining neuronal integrity.

In summary, SMC-mediated reduction of *4R MAPT* transcript expression lowers expression of 4R-tau protein in S305N and in P301L neurons, ultimately influencing overall tau accumulation. As a result, we also observed a decrease in accumulation of phosphorylated tau species pS202/T205 and pS396, both markers strongly associated with tau pathology. Notably, the most pronounced reduction occurred for oligomeric pTau-S396 (>250 kDa) with reduced SDS solubility. Our findings suggest that reducing the disease-causing mutant 4R tau expression (P301L) or correcting 4R/3R splicing (S305N) induces a global shift in tau accumulation, leading to decreased pTau oligomerization. Based on our analysis, we ranked our lead compounds by tau reduction effectiveness, with PTC-775 exhibiting the strongest and most consistent effect, followed by PTC-700 and PTC-553.

### SMCs reduce insoluble tau and mitigate stress vulnerability in tauopathy neurons

A key factor in the evaluation of the therapeutic potential of new small molecules is their ability to target and mitigate the most relevant disease hallmarks, thereby supporting their biological and clinical relevance. In tauopathy neuronal models, one such hallmark is the accumulation of pTau protein species with reduced detergent solubility (*65–68*). To assess the effect of SMCs on tau solubility, we focused on the top two candidates, PTC-700 and PTC-775, using tau-P301L neurons as a representative tauopathy model that captures the accumulation of insoluble tau species (*35, 36, 63*). In P301L iPSC-derived neurons, some tau species with reduced SDS solubility appeared as high MW bands on SDS-PAGE, detected with the pTau-S396 antibody (Fig. 3, Fig. S4). We found that both PTC-700 and PTC-775 effectively reduced the levels of these insoluble species. To further characterize tau solubility changes, we performed detergent fractionation of neuronal lysates into soluble and insoluble protein fractions (Fig. 4A). As in the previous analyses, we treated neurons for 96h (1 μM SMC) or 3 weeks (100 nM SMC) with the concentration corresponding to ∼EC50 for each treatment regimen (see Fig. 3, Fig. S4). After treatment, cells were lysed and subjected to protein fractionation based on differential solubility in Triton X-100 and SDS detergents (Fig. 4A) (*69, 70*). Western blot analysis of soluble (S) and insoluble-pellet (P) protein fractions, in SMC-treated neurons relative to vehicle controls, enabled us to assess the effect of each SMC on total tau (TAU5) and pTau-S396 soluble and insoluble species. Treatment with PTC-700 (Fig. 4B, C) and PTC-775 (Fig. 4D, E) generally resulted in a reduction of both total tau and pTau species across fractions. PTC-700 induced a similar reduction in both soluble and insoluble tau and pTau-S396 after both 96h (Fig. 4B) and 3 weeks of treatment (Fig. 4C), with the reduction in insoluble pTau-S396 being somewhat more pronounced. Prolonged treatment resulted in a similar decrease in insoluble tau species but achieved at lower concentrations, with total tau decreasing by approximately 40% and pTau-S396 by approximately 70% (Fig. 4C). PTC- 775 demonstrated a stronger effect after 3 weeks of treatment and at a lower concentration (Fig. 4E vs. D), reducing insoluble total tau by approximately 90% and insoluble pTau-S396 by ≥ 70%. At these concentrations, the SMCs effectively reduced insoluble pTau species, which are hypothesized to be the most aggregation-prone and toxic species driving disease progression. The persistence of residual insoluble pTau could be due to the maximum tau reduction not being achieved at EC50 and some mutant 4R tau remaining in the neurons, and/or additional factors in FTD patient-derived neurons contributing to this pool of misfolded tau.

**Figure 4.**
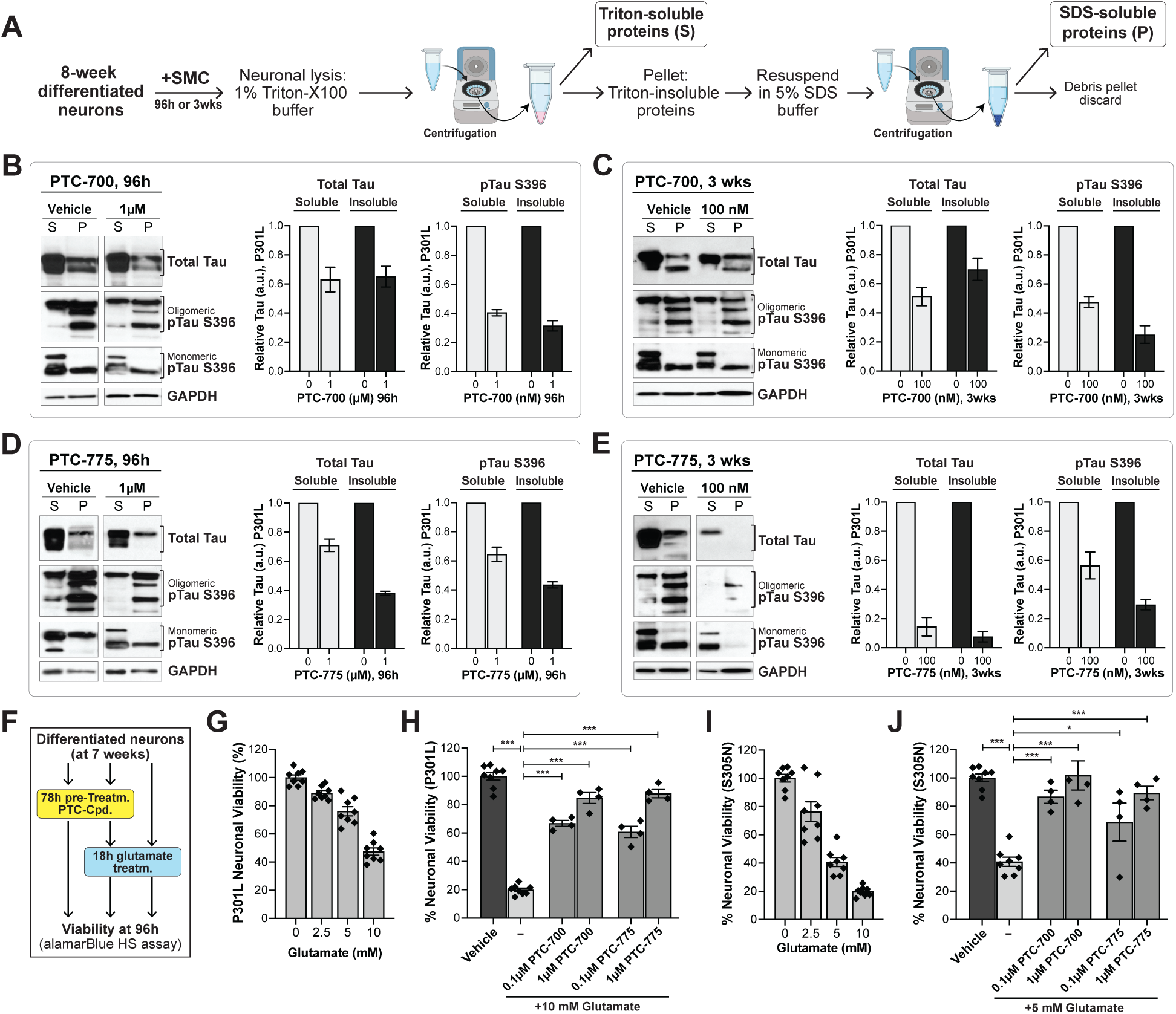
SMC treatment reduces insoluble tau and rescues FTD neuronal stress vulnerability. **(A)** Schematic of the assay employed to test protein solubility. Neuronal lysates are fractionated based on proteins’ differential solubility to TritonX-100 and high % SDS detergents and analyzed as two protein fractions designated soluble (S) or insoluble (P). **(B-E)** Western blot and densitometry analysis of soluble (S) and insoluble (P) tau in P301L neurons treated with PTC-700 and PTC-775 for 96h (B, D) or for 3 weeks (C, E). Graph bars represent mean densitometry normalized to GAPDH (S) and relative to DMSO-treated neurons (0 μM) ± SD, with *N* = 2 biological replicates. **(F)** Assay to measure stress vulnerability in neurons pre-treated with SMC or DMSO (control). **(G)** Eight-week differentiated tau-P301L and **(I)** tau- S305N neurons show loss of cell viability when treated with increasing concentrations of glutamate for 18h. Individual data points are indicated and graph bars represent mean neuronal viability (%) relative to vehicle (0 μM = 100%) treated neurons ± SEM (*N* = 4, with 2 technical replicates included per assay). When pre-treated with PTC-700 or PTC-775 for 78h (total 96h of treatment), 8-week differentiated and glutamate-stressed **(H)** P301L and **(I)** S305N neurons show improved cell viability relative to glutamate-only treated neurons (*light grey bar*). Individual data points are indicated and graph bars represent mean neuronal viability (%) relative to DMSO-treated neurons ± SEM (*N* = 4). Statistical analysis was performed using a two-tailed, paired Student’s t-test. Significance thresholds: \**p* ≤ 0.05, and \*\*\**p* ≤ 0.001.

To demonstrate the therapeutic relevance of SMC-mediated 4R tau reduction and the resulting decrease in disease-associated tau species, including hyperphosphorylated and insoluble tau, we evaluated the ability of SMCs to mitigate tau toxicity. In FTD patient-derived neurons, aberrant accumulation of tau is associated with increased neuronal vulnerability to stress (*36, 64*)*, i.e.* increased cell death in the presence of specific stressors that, in the same conditions, do not affect non-disease neurons. One of the stressors that reveals this phenotype is a high dose of the excitatory neurotransmitter glutamate. Glutamate promotes significant concentration-dependent loss of viability in P301L (Fig. 4G) and S305N (Fig. 4I) neurons, but not in non-mutant neurons (Fig. S6A), revealing mutant tau-dependent neuronal vulnerability. These findings align with our previous demonstration that CRISPR-mediated *MAPT* knockout rescues vulnerability to stress, demonstrating tau-mediated toxicity in these neuronal models (*64*). In this study, we investigated if PTC-700 and PTC-775, by reducing 4R tau levels, confer protection against stress vulnerability using the stress vulnerability assay (Fig. 4F). In the P301L model (Fig. 4H), neurons pre-treated with vehicle (78h) followed by 10 mM glutamate (18h) exhibited a viability loss to 20% of the vehicle control. When pre-treated with PTC-700 or PTC-775, we observed a rescue in cell viability to 60–90% of the vehicle control, with the higher SMC concentration (1 μM vs. 100 nM) producing a proportionally greater protective effect (Fig. 4H). Similarly, S305N neurons pre-treated with vehicle (78h) followed by 5 mM glutamate (18h), exhibited a viability loss to 40% of the vehicle control (Fig. 4J). However, when pre-treated with PTC-700 or PTC-775, we observed a rescue in cell viability to 80–100% of the vehicle control, with the higher SMC concentration (1 μM vs. 100 nM) providing a stronger protective effect (Fig. 4J). Neurons treated with vehicle or SMC alone showed no changes in viability (Fig. S6B, C).

Taken together, these findings demonstrate that PTC-700 and PTC-775 effectively reduce 4R tau expression and mitigate the 4R-tau-driven accumulation of total tau and insoluble pTau species in P301L and S305N neurons. Notably, the reduction in 4R tau and its associated phenotypes translates into a significant decrease in tau-associated neuronal toxicity, underscoring the potential therapeutic value of our SMCs for tauopathies.

### Enhanced efficacy of PTC-003 rescuing tauopathy phenotypes in patient iPSC-derived neurons

Building on the promising results with our lead SMCs in mutant tau iPSC-derived neurons, we examined the *in vivo* pharmacokinetic (PK) properties of our SMC collection and identified close analogs with improved CNS penetrance to enable pharmacodynamic (PD) studies. Among the compounds identified, SMC PTC-003 demonstrated superior CNS biodistribution and was consequently selected for *in vivo* testing (Fig. 1C). PTC-003 was quite potent in the *in vitro* minigene splicing assay, inducing a dose-dependent increase in *3R MAPT* (EC2X = 5.2 nM, Fig. 5A) and corresponding decrease in *4R MAPT* (Fig. 5A: *bottom gel panel*). Consistent with this result, in the MSD protein assay, we observed a dose-dependent increase in 3R tau (Fig. 5B). The threshold concentration for a near-maximal effect (EC1.3X) was 7.4 nM, representing an improvement over previous compounds, including PTC-775 (EC1.3X = 8.9 nM, Fig. 1J).

**Figure 5.**
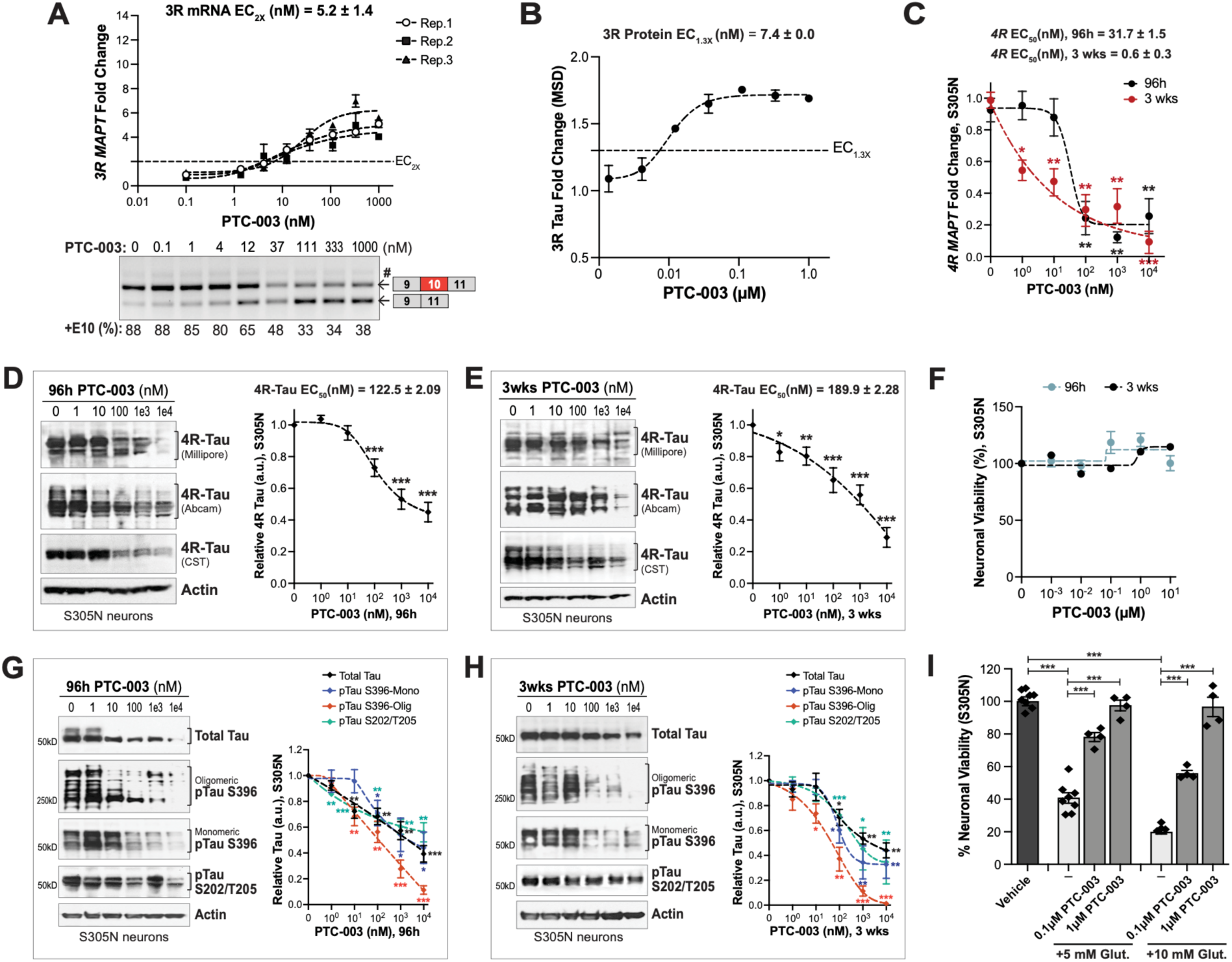
PTC-003 lowers *4R MAPT* expression and rescues tauopathy phenotypes in the S305N iPSC-neuronal model. **(A)** *MAPT* minigene assay in HEK293T cells treated with PTC-003 for 24h. RT- qPCR analysis to measure exon 10 skipping and the corresponding increase in *3R MAPT* (EC_2X_ indicated). Data points represent fold-change relative to DMSO-treated cells ± SEM, with *N* = 3. Representative RT- PCR analysis of *MAPT* splicing is shown below, demonstrating dose-dependent increase in *3R MAPT* (– E10, lower band) and a corresponding decrease in *4R MAPT* (+E10, upper band). A faint upper band (#) is also visible, which shares the same nucleotide sequence as the *4R MAPT* band and likely represents a secondary structure of the PCR product. **(B)** MSD assay reveals compound-mediated increase in 3R tau protein in U-87 glioma cells. Data points represent mean fold-change relative to DMSO-treated cells ± SEM with *N* = 3 (EC_1.3X_ shown). **(C)** RT-qPCR analysis of SMC dose curves for *4R MAPT* in S305N neurons following 96h or 3 weeks treatment (EC_50_ indicated). Data points represent mean fold change relative to DMSO-treated neurons (normalized to *GAPDH*) ± SEM, with *N* = 2 (3 technical replicates included per assay). **(D, E)** Western blot analysis of SMC effect on 4R tau of S305N neurons after 96h (D) or 3 weeks (E) of treatment, with representative blots shown. Densitometry analysis of 4R tau, relative to DMSO-treated (0 μM) neurons (normalized to actin). EC_50_ is indicated. Data points represent mean densitometry ± SEM, with *N* = 3. **(F)** Viability dose curves for 8-week differentiated S305N neurons treated with PTC-003 for 96h or 3 weeks. Data points represent mean viability (%) relative to DMSO-treated neurons (100% viability) ± SEM, with *N* = 3. **(G, H)** PTC-003 dose curves for total tau (TAU5 antibody), pTau-S396 (high MW–oligomeric and monomeric pTau), and pTau-S202/T205 (AT8 antibody) after 96h (G) or 3 weeks (H) of treatment, by western blot and densitometry analysis in S305N neurons. Representative blots are shown. Data points represent mean values relative to DMSO-treated neurons ± SEM, with *N* = 3. Statistical analysis was performed using a two-tailed, paired Student’s t-test for each compound concentration vs. vehicle (0 μM). **(I)** When pre-treated with PTC-003 (78h) and stressed with glutamate (5 mM or 10 mM), S305N neurons show improved neuronal viability relative to glutamate-only treated neurons (*light grey bars*). Individual data points are indicated and graph bars represent mean neuronal viability (%) relative to DMSO ± SEM (*N* = 4). Statistical analysis was performed using a two- tailed, paired Student’s t-test. Significance threshold \*\*\**p* ≤ 0.001.

Before advancing to *in vivo* studies, we evaluated PTC-003 in tau-S305N neurons. This tauopathy neuronal model was specifically selected because, it carries a splicing *MAPT* mutation that upregulates 4R tau expression, a similar type of mutation as the one expressed in the animal model we have selected for *in vivo* studies. As in the earlier work, we treated S305N neurons for either 96h or 3 weeks (Fig. 2A) and measured *4R MAPT* mRNA levels using RT-qPCR (Fig. 5C). Both treatment regimens resulted in a significant dose-dependent reduction in *4R MAPT*, with the 3-week treatment producing a lower EC50 (0.6 nM) compared to the 96h treatment (EC50 = 31.7 nM, Fig. 5C). PTC-003 exhibited a slightly stronger effect on *4R MAPT* transcript reduction than all other tested SMCs. At the protein level, we observed a concentration-dependent reduction in 4R tau, with comparable EC50 magnitudes (123-190 nM) for both treatment regimens (Fig. 5D, E). However, the 3-week treatment led to a more pronounced reduction of tau, reaching nearly 80%. Importantly, PTC-003 treatment did not compromise neuronal viability; instead, we observed a modest increase in viability at concentrations ≥ 100 nM (Fig. 5F).

To evaluate the effect of PTC-003 on tauopathy phenotypes, we assessed treatment impact on the accumulation of total tau, pTau-S396 (monomeric and oligomeric species), and pTau- S202/T205 (AT8) following 96h (Fig. 5G) or 3 weeks (Fig. 5H) of treatment. After 96h, we observed a clear dose-dependent reduction across all markers, with most tau species decreasing by approximately 60% and oligomeric pTau-S396 exhibiting an impressive > 90% reduction at the 10 μM dose (Fig. 5G). The 3-week treatment further amplified these effects, achieving over 90% reduction of oligomeric pTau-S396 at 1 μM SMC (Fig. 5H), demonstrating the increased efficacy of prolonged treatment. To determine whether PTC-003 correction of 4R/3R splice balance and reduction of tau accumulation mitigates neuronal toxicity, we performed the stress vulnerability assay (Fig. 4F). At the beginning of the eighth week of differentiation, S305N neurons were pre-treated with either 0.1 μM or 1 μM PTC-003, or vehicle (DMSO) alone, followed by an 18h glutamate stress (5 mM or 10 mM). In neurons pre-treated with vehicle, 5 mM glutamate reduced viability to 40% of vehicle-alone control neurons, while 10 mM glutamate further decreased viability to 20% (Fig. 5I, *lightest bars*). Notably, pre-treatment with PTC-003 resulted in a dramatic rescue of viability, restoring it to 80–100% in 5 mM glutamate-stressed cells (Fig. 5I, *middle bars*) and to 60–90% in neurons exposed to 10 mM glutamate (Fig. 5I, *rightmost bars*). We also detected that the rescue effect was more pronounced at the higher PTC-003 concentration (1 μM vs. 0.1 μM), further supporting its dose-dependent protective effect against tau toxicity.

These findings demonstrate that PTC-003 effectively reduces tau accumulation, particularly pTau oligomeric species, while also mitigating tau-mediated toxicity. Neuronal pre-treatment with PTC-003 also significantly protected neurons against stress conditions, highlighting its therapeutic potential.

### PTC-003 ameliorates tau pathology in the human tau-N279K gene-replacement mouse model

To evaluate the *in vivo* efficacy and therapeutic relevance of the SMC PTC-003, we utilized a new FTD mouse model (*61*). This model was generated using a gene-replacement strategy (*MAPT*- GR*N279K), in which the mouse *Mapt* gene is fully replaced by the human *MAPT* gene, resulting in expression of human tau-N279K within the H1 haplotype, on a murine *tau*-null background (*61*). To assess genotype-dependent expression of human tau, we analyzed the levels of *MAPT* isoforms and found that homozygous tau-N279K/N279K mice exhibited increased human *4R MAPT* and reduced *3R MAPT* mRNA levels as early as one month of age (Fig. 6A). The *4R/3R MAPT* ratio was higher in homozygous mice compared to N279K/WT heterozygous mice (Fig. 6A), with both groups displaying significantly elevated *4R MAPT* relative to wild type (WT/WT) controls (Fig. 6A). These results indicate that both heterozygous and homozygous tau-N279K animals are suitable for testing the *in vivo 4R MAPT*-reducing effects of SMCs.

**Figure 6.**
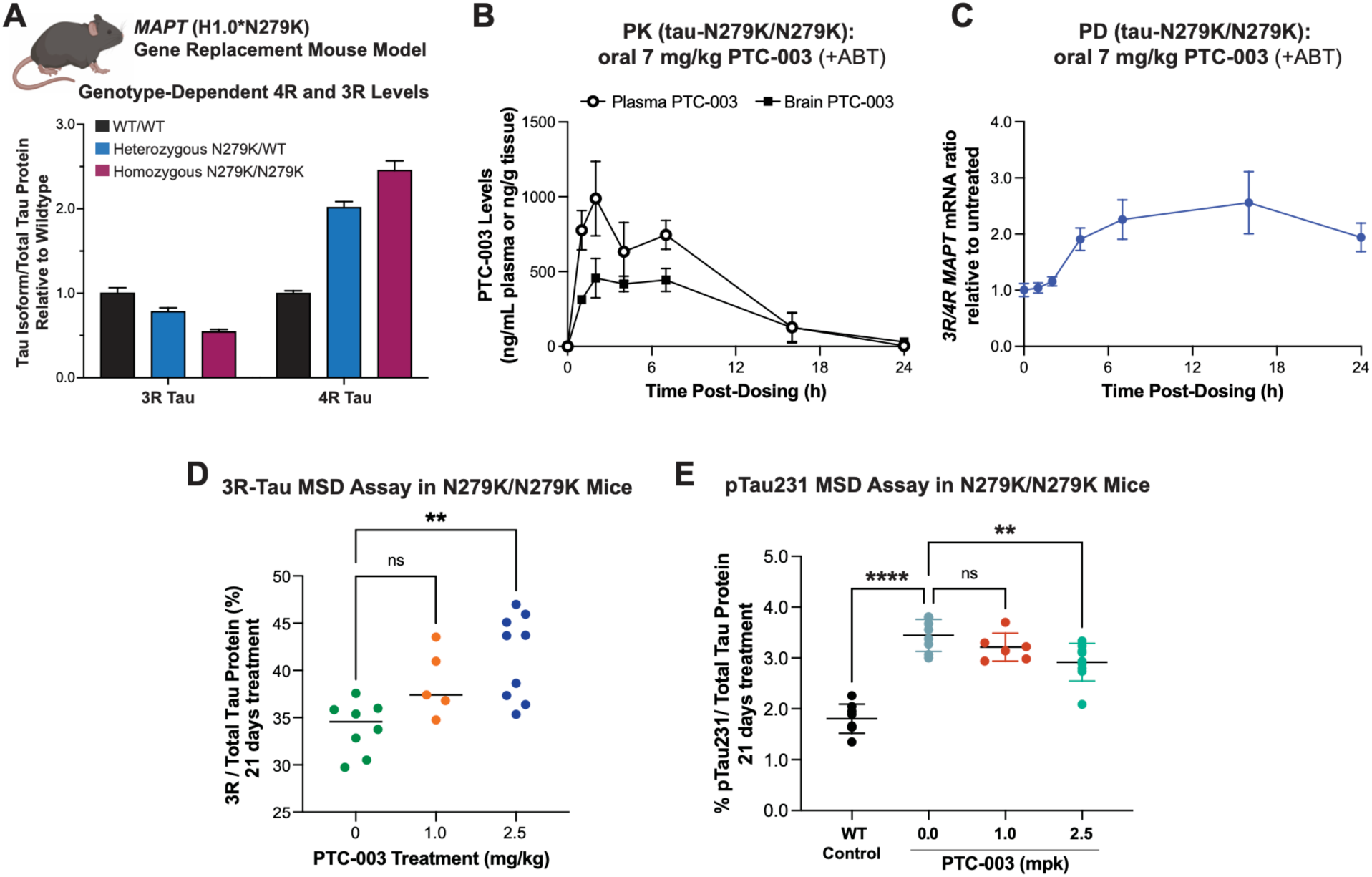
Oral administration of PTC-003 corrects *MAPT* splicing and reduces pTau231 in the brain of a human GR tau-N279K mouse model. **(A)** Genotype-dependent differences in relative levels of 4R and 3R tau in WT/WT, heterozygous N279K/WT, and homozygous N279K/N279K tau mice. Data bars show the mean values from 5 mice ± SEM. **(B)** Pharmacokinetics (PK) and **(C)** pharmacodynamics (PD) measurements after a single dose of PTC-003 administered to tau-N279K/N279K mice. PTC-003 was dosed at 7 mg/kg via oral gavage in the presence of ABT (50 mg/kg). Data points represent the mean of 3 mice/group ± SEM. In (C) the 3R/4R ratio (shown as 1 at 0h) was normalized to untreated mice, otherwise *3R/4R MAPT* <1 in tau-N279K homozygous animals, from (A). **(D)** Increase in brain 3R tau in N279K homozygous mice after daily oral dosing with PTC-003 (+ABT) for 21 days, measured by the MSD assay. Data points represent individual animals. **(E)** Decrease in brain levels of pTau231 in N279K homozygous mice after daily oral dosing with PTC-003 for 21 days. Data points represent individual animals. Shown for comparison are levels in WT mice. Statistical analysis was performed with one-way ANOVA and Dunnett’s multiple comparisons test. Significance thresholds: non-significant (n/s) *p* > 0.01, or \*\**p <* 0.001 and \*\*\*\**p <* 0.00001.

Despite favorable CNS biodistribution, the pharmacokinetics of PTC-003 following oral administration were suboptimal due to rapid liver clearance, with a significant decline in compound levels in both plasma and brain observed at ≤ 7h post-dosing (Fig. S7A, B). To address this limitation, we administered the pan-cytochrome P450 inhibitor 1-aminobenzotriazole (ABT) (*71*) 2h prior to dosing PTC-003 via oral gavage. This increased PTC-003 exposure in plasma and brain (Fig. S7A, B), resulting in brain concentrations predicted to be above the *in vitro* EC50 (Fig. 5C-D). We then conducted two *in vivo* studies in the homozygous tau-N279K/N279K mouse. In the first study, executed to measure the SMC effect on *MAPT* transcript levels, 2-month-old mice were orally treated with a single dose of 7 mg/kg PTC-003 combined with 50 mg/kg ABT (Fig. 6B, C). Mice were euthanized at 1, 2-, 4-, 7-, 16-, and 24-hours post-dosing, and plasma and brain tissues were collected for analysis of compound levels (Fig. 6B), as well as *3R* and *4R MAPT* transcript levels (Fig. 6C). This analysis confirmed that co-treatment with ABT enhanced brain exposure of the compound in mice, enabling detection of PTC-003 in brain tissue (Fig. 6B). Consequently, PTC-003 induced a dose-dependent increase in the *3R/4R MAPT* splice ratio, with a peak observed 16h post-dosing (Fig. 6C).

We conducted a follow-up study to determine whether the increase in *3R* mRNA transcript levels corresponded to an increase in 3R tau protein levels in the brain. Two-month-old homozygous tau-N279K/N279K mice were treated daily via oral gavage with PTC-003 at doses of 1 mg/kg and 2.5 mg/kg for 21 days, in the presence of ABT. After treatment, the mice were euthanized, and brain tissues were processed for analysis of 3R tau, total tau (Fig. 6D), and pTau- T231 (Fig. 6E) protein levels (*72, 73*). Using our MSD protein assay, we observed a dose-dependent increase in the 3R tau/total tau ratio following PTC-003 treatment, which reached statistical significance at the highest dose of 2.5 mg/kg (Fig. 6D). Moreover, homozygous N279K mice displayed a statistically significant increase in pTau-T231 levels in the brain compared to control WT animals (Fig. 6E). This pTau marker, referred to as pTau231, is a well-studied pathological biomarker in tauopathies, including FTD and Alzheimer’s disease, reporting on early tau hyperphosphorylation and conformational changes (*72, 73*). Importantly, PTC-003 reduced pTau231 in a dose-dependent manner, with a significant reduction observed with the 2.5 mg/kg treatment (Fig. 6E).

Overall, these findings in the human gene-replacement tau-N279K mouse model provide strong *in vivo* evidence of the therapeutic potential of our lead SMC PTC-003 in promoting *MAPT* exon 10 skipping, increasing *3R MAPT* transcript and protein levels in the brain as a consequence of reduced *4R MAPT*, and mitigating tauopathy-relevant phenotypes, including pTau231 accumulation.

## DISCUSSION

Tauopathies are neurodegenerative diseases strongly linked to the pathological accumulation of tau protein in affected brain regions. In some familial forms of the disease, mutations in *MAPT* exon 10 and adjacent intronic regions can either disrupt splicing, altering the relative expression of 3R and 4R tau isoforms, or confer gain-of-function properties to the 4R tau protein that enhance its propensity to misfold and accumulate (Fig. 1A, B). In this study, we investigated three mutations relevant to a significant proportion of FTD patients (Fig. 2A): tau-P301L, a gain-of-function mutation that produces mutant 4R tau isoforms with a high propensity for aggregation (*16*); and the splicing mutations S305N and N279K, both of which increase the 4R/3R splice ratio by promoting inclusion of *MAPT* exon 10, thereby enhancing 4R MAPT transcript and protein expression (*74*). Given the pathological consequences of these mutations, a therapeutic strategy that promotes *MAPT* exon 10 exclusion may offer a clinical benefit.

Currently, no disease-modifying therapies exist for tauopathies, and only a few experimental tau-targeting drugs are in clinical trials (*19–36*). Genetic approaches to modulate *MAPT* splicing, such as ASOs (*20, 21, 37, 38*), have shown promise. However, even the most advanced ASO-based therapies face critical challenges, including poor brain penetration, limited cellular uptake, immune-related toxicity, and the need for invasive administration routes (*40–43*). Given the inherent hurdles in developing therapies for neurodegenerative diseases, exploring multiple therapeutic strategies is essential. In this context, small-molecule SMCs are a promising alternative or complementary therapeutic approach (*45–47*). Small-molecule SMCs offer significant advantages as therapeutics for neurological disorders, including oral bioavailability, broad brain penetration, and the ability to be administered systemically without invasive procedures. In this study, we evaluated the efficacy and therapeutic potential of a newly developed class of SMCs in two types of models: FTD patient iPSC-derived neuronal models, which offer a physiological, cellular, and genomic relevant platform for studying *MAPT* splicing modulation; as well as a new gene replacement (GR) tauopathy mouse model. Our SMCs were designed and screened for their ability to promote *MAPT* exon 10 skipping. We demonstrated that SMC treatment reduces 4R tau expression, leading to a downstream decrease in accumulation of total tau and pTau, including insoluble oligomeric proteoforms. The lead compounds, PTC-700, PTC- 775 and PTC-003, demonstrated efficacy in iPSC-derived neurons at nanomolar concentrations, exerting measurable effects after a single 96h treatment (Fig. 2-3). Their impact was further enhanced following a 3-week treatment (Fig. 2, S4), which lowered EC50 values and led to a more potent reduction of total tau and pTau at the higher concentrations. Notably, both treatment regimens were well tolerated (Fig. 3A-C, S4A-C), and at the highest SMC doses, PTC-775 (Fig. S4C) and PTC-003 (Fig. 5F) showed a modest improvement in neuronal viability. These findings highlight the exceptional potency and efficacy of our novel small-molecule regulators of *MAPT* exon 10 splicing, demonstrating a strong ability to reduce 4R expression and to mitigate 4R-tau-dependent tau accumulation in FTD patient iPSC-derived neurons. Additionally, we evaluated the impact of our SMCs on key tauopathy phenotypes, focusing on their effect on detergent-insoluble, aggregation-prone tau and tau-driven neurotoxicity, assessed by the neuronal stress vulnerability assay. Our findings demonstrate that lead SMCs significantly reduced both soluble and insoluble tau in P301L neurons, with the effect being further amplified by a prolonged 3-week treatment, leading to over 80% reduction in insoluble tau (Fig. 4D, E). Consistent with these phenotypic changes, lead SMCs also mitigated the vulnerability of P301L and S305N neurons to glutamate-induced excitotoxicity (Fig. 4G-J). Under basal conditions, P301L and S305N neurons exhibit distinct sensitivities to glutamate, consistent with their different mutation-driven pathologies (*12, 17*). Tau-S305N neurons, which accumulate higher levels of 4R tau, show increased susceptibility to glutamate toxicity, as indicated by reduced neuronal viability at high glutamate concentrations (Fig. 4G vs. 4I). Importantly, pre-treatment with the lead SMCs, PTC-700, PTC-775 and PTC- 003, significantly rescued neuronal viability to above 80% compared to vehicle-treated neurons (100% viability, Fig. 4H, 4J, 5I).

Building on the promising results in patient-derived iPSC neurons, and focusing on PTC- 003, a close analog of PTC-700 with optimized pharmacokinetic properties, we then evaluated the *in vivo* efficacy of PTC-003 in the human GR-tauopathy mouse model expressing tau-N279K (*61*). We selected this model because it was generated by replacing the murine *Mapt* locus with the full human *MAPT* gene, including the intronic regions, allowing us to test the modulation of *MAPT* exon 10 splicing, while faithfully recapitulating specific human disease mechanisms (*61*). PTC-003 effectively modulated *MAPT* exon 10 splicing, leading to an increase in 3R/4R tau ratio in the brain of homozygous tau-N279K/N279K mice (Fig. 6B, C). Most importantly, treatment with PTC-003 significantly reduced pTau231 accumulation (Fig. 6E), underscoring the remarkable therapeutic potential of this novel class of small molecules as an oral treatment for tauopathies.

Collectively, our findings provide compelling evidence that splicing modulators represent a promising therapeutic strategy for tauopathies. We demonstrated the potency and efficacy of our SMCs in significantly reducing 4R tau and mitigating 4R-dependent tauopathy phenotypes in patient-derived neuronal models and in a human GR tauopathy mouse model. While these studies focused on genetic forms of disease, this innovative approach may also benefit sporadic tauopathies characterized by abnormal 4R tau accumulation. Therefore, these results lay the foundation for future preclinical and clinical development of small-molecule splicing modulators as a viable therapeutic strategy for FTD and related tauopathies.

## MATERIALS AND METHODS

### Study design

This study was designed to evaluate the therapeutic potential of small-molecule SMCs targeting *MAPT* exon 10 splicing in 4R tauopathies. We tested the hypothesis that reducing 4R tau expression through *MAPT* splicing correction would mitigate tau pathology and neuronal toxicity. To this end, we employed patient-derived neuronal models carrying the tau P301L (gain-of-function) and S305N (splicing) mutations, as well as a human gene-replacement mouse model (*MAPT*-GR*N279K) expressing human tau under the H1 haplotype on a murine *tau*-null background. Neurons were treated with SMCs for either 96h or 3 weeks *in vitro*, while animals received daily oral administration for 24h or 21 days. DMSO-treated cultures and vehicle-treated animals served as controls. Outcome measures included quantification of 4R tau (transcript and protein), total tau, and phospho-tau species by RT-PCR and western blot, along with assessments of neuronal viability and tau-mediated toxicity. *In vivo* efficacy was evaluated by biochemical analysis of brain tissue and plasma levels of *MAPT* transcripts and pTau protein. All experiments were performed with biological replicates (*N* ≥ 3), with treatment assignments randomized and analyses conducted blinded to condition. Statistical analyses are detailed in the figure legends and Methods.

### Compounds synthesis and quality control

#### Synthesis

Small-molecule splicing modulators PTC-700 and PTC-553 were synthesized as described in published methods (WO 2021/118929 A1, “*Compounds for Treating Familial Dysautonomia*” and WO 2020/167628 A1, “*Thioeno[3,2-B] Pyridine-7-Amine Compounds for Treating Familial Dysautonomia*”). PTC-775 and PTC-003 are close analogs of PTC-700. Purities of all compounds were found to be >95% as determined by reverse-phase LC-MS and ^1^HNMR. ^1^HNMR spectra were recorded on a Bruker Avance III spectrometer (500 MHz, ^1^H) equipped with a 5 mm Bruker PA BBO probe. Spectra are referenced to the residual solvent signal: perdeuterated methanol (CD3OD, δ 3.31 ppm, ^1^H). Chemical shifts are reported in ppm (δ); multiplicities are indicated by s (singlet), d (doublet), t (triplet), q (quartet), m (multiplet), and in combination. Coupling constants, *J*, are reported in Hz. Spectra were acquired at a temperature of 298 K (25°C). Liquid Chromatography-Mass Spectrometry (LC-MS) samples were run on a Waters Acquity UPLC I-class system with Flow Through Needle sample manager. The gradient was created using a binary solvent pump. Separations were achieved with a Waters Acquity HSS C18 1.8 μm MVK column with Vanguard, held constant at 45°C. Analytes were detected with a photodiode array eλ detector scanning a range of 210-400 nm, focused on 254 nm, and a QDa detector using a mass range of 150-1000 m/z. Chromatography results are reported as R*f* (retention factor in minutes (min)); spectral results are reported as (mass, intensity).

*2-[(2S)-2-Aminopropyl]-5-chloro-3-methyl-N-[(1,3-thiazol-2-yl)methyl]thieno[3,2- b]pyridin-7-amine* (PTC-700). ^1^HNMR (500 MHz, CD3OD, 298 K): δ 7.77 (d, *J* = 3.2 Hz, 1H), 7.53 (d, *J* = 3.2 Hz, 1H), 6.44 (s, 1H), 4.87 (s, 2H), 3.25 (h, *J* = 6.8 Hz, 1H), 2.98 (dq, *J* = 15.5, 7.2 Hz, 2H), 2.33 (s, 3H), 1.16 (d, *J* = 6.3 Hz, 3H); LCMS (ESI^+^, TOF): 177.2 (20), 353.2 (100), 355.2 (40), 356.1 (5).

*6-[(2S)-2-aminopropyl]-7-bromo-2-chloro-N-[(thiophen-2-yl)methyl]thieno[3,2- d]pyrimidin-4-amine* (PTC-553). ^1^HNMR (500 MHz, CD3OD, 298 K): δ 8.54 (s, 1H), 7.28 (dd, *J* = 5.2, 1.2 Hz, 1H), 7.09 (dd, *J* = 3.4, 1.2 Hz, 1H), 6.95 (dd, *J* = 5.1, 3.4 Hz, 1H), 4.92 (s, 2H), 3.65 (dq, *J* = 13.0, 6.7 Hz, 1H), 3.34 – 3.20 (m, 2H), 1.33 (d, *J* = 6.6 Hz, 3H); LCMS (ESI^+^, TOF): 415.1 (5), 417.1 (65), 419.1 (100), 421.1 (30), 441.0 (5).

### HEK293T-MAPT cells and compound treatment

HEK293T (ATCC–HEK293T Human Kidney) cells carrying a stably-integrated minigene for mutant *MAPT* (splicing mutation IVS10+16) were cultured as previously described (*62*). Briefly, cells were cultured in Dulbecco’s modified Eagle’s medium (D-MEM, Gibco 11995-065) supplemented with 10% fetal bovine serum (FBS, Sigma 12306C) and 1% penicillin/streptomycin (Pen/Strep, Corning 30-009-CI). Compounds PTC-003, PTC-553, PTC-700 and PTC-775 were initially dissolved in 100% DMSO (Sigma D8418) to yield 10 mM stock solutions. Working solutions (200X) for different concentrations of the compounds were prepared by further diluting the stock solutions in 100% DMSO. The final DMSO concentration in the compound-treated or vehicle-treated cells was 0.5% (v/v). For compound treatment, HEK293T cells were plated at a density of ∼74,000 cells/cm^2^ in 24-well plates, and after 24h the medium was replaced with 500 μL of complete DMEM (see above) containing the appropriate concentration of a specific compound or DMSO. After 24h of incubation, cells were collected for total RNA analysis.

### HEK293T-MAPT cells RNA isolation, RT-PCR and RT-qPCR analysis

PBS-washed cells were collected into a pellet by centrifugation (3,000 *xg* for 5 min) and RNA was extracted with the QIAzol Lysis Reagent (Qiagen 79306) following the manufacturer’s instructions. Total RNA yields were determined using a Nanodrop ND-1000 spectrophotometer. Reverse transcription and cDNA synthesis were performed using 0.5-1 µg of total RNA, Random Primers (Promega C1181), Oligo(dT)15 Primer (Promega C1101), and Superscript III reverse transcriptase (ThermoFisher Scientific 18080093) according to the manufacturer’s protocol. cDNA was used to setup PCR reactions in a 20-25 µL volume, with GoTaq Green Master mix (Promega MT123). To detect both 4R and 3R isoforms we used MAPT_ex10_F forward primer 5’-CCCAAGTCGCCGTCTTCC-3’ and MAPT_ex11_R reverse primer 5’- TGGTTTATGATGGATGTTGCCT-3’. The PCR reaction was: 95°C for 3 min; 34 cycles of [95°C for 30 s, 55°C for 30 s, 72°C for 30 s]; final extension at 72°C for 5 min. PCR products were resolved on a 1.8% (w/v) agarose gel in TAE buffer and visualized by ethidium bromide staining. The relative amount of *4R* and *3R MAPT* isoforms were determined using the integrated density value (IDV) for each transcript band, assessed using Alpha 2000TM Image Analyzer and quantified by ImageJ software. Data analysis was done in Microsoft Excel and values were plotted in GraphPad Prism 10.

The mRNA levels of *3R MAPT* and *GAPDH* were quantified by reverse-transcription quantitative real-time PCR (RT-qPCR) analysis using The LightCycler-480 Instrument (Roche) and Taqman-based probes (ThermoFisher Scientific). The reactions were setup with a cDNA equivalent of 25 ng of starting RNA in a 20 μL reaction volume. To amplify the *3R MAPT* we utilized: Taq_3R_qPCR forward 5’-AGGCGGGAAGGTGCAAATA-3’; Taq_3R_qPCR reverse 5’-CTGGTTTATGATGGATGTTGCCT-3’; Taq_3R_qPCR probe 5’-TCTACAAACCAGTTGACCTGAGCAAGGTGACC-3’. To amplify *GAPDH* we utilized Taq_hGADPH_qPCR forward 5’-CAACGGATTTGGTCGTATTGG-3’; Taq_hGADPH_qPCR reverse 5’-TGATGGCAACAATATCCACTTTACC-3’; Taq_hGADPH_qPCR probe 5’- CGCCTGGTCACCAGGGCTGCT-3’. The forward and reverse primers were used at a final concentration of 0.4 μM in the PCR reaction. The probes were used at a final concentration of 0.15 μM. RT-qPCR was carried out with the following settings: step 1 at 95°C for 10 min; step 2 at 95°C for 10 s; step 2 at 60°C for 30 s; step 3 at 72°C for 1 min; steps 2 to 3 were repeated for 45 cycles. Data were analyzed with the The LightCycler 480 software (Roche) and normalized to *GAPDH*. Fold changes were calculated relative to vehicle (DMSO) treated sample. Values were plotted in GraphPad Prism 10.

### Small-molecule screening in U87-MG cells

U87-MG cells were maintained in MEM media supplemented (MEM^+^) with 10% fetal bovine serum, 1% penicillin/streptomycin, 1% non-essential amino acids, and 1% sodium pyruvate (all Gibco/Life Technologies). For compound screening in 96-well Nunc plates (ThermoFisher Scientific), U87 cells were plated at 15,000 cells/well in 200 μL of MEM^+^ media pre-treated with compounds at a 7-point concentration range. The treatment plates were incubated at 37°C and 5% CO2 for 96h. After 96h, media was aspirated and 120 μL of MSD Tris lysis buffer (Mesoscale Discovery), supplemented with 1x Halt protease inhibitor cocktail (ThermoFisher Scientific), was added per well to promote cell lysis. Plates were incubated at room temperature for 15 minutes on an orbital shaker. Lysates were posteriorly analyzed with the MSD assay.

### Meso Scale Discovery (MSD) immunoassay for 3R tau and total tau quantification in cells

MSD multi-array plates (Mesoscale Discovery, L15XA-3) were coated with total tau monoclonal antibody (Cell Signaling Technology, clone D5D8N, rabbit). Briefly, the antibody was diluted to 0.5 µg/mL in 1X PBS (Gibco/Life technologies) and 50 µL/well was added to the MSD plate.

Plates were sealed, centrifugated (short spin), and placed at 4°C overnight. The detection antibody 3R tau (Millipore-Sigma, clone 8E6/C11, mouse) was diluted 1:250 in 1% Blocker A (Mesoscale Discovery R93AA-1) + 0.05% Tween20. Goat anti-mouse SULFO-tagged antibody (Mesoscale Discovery R32AC) was diluted 1:500 in 1% Blocker A + 0.05% Tween20. Plates were sealed, centrifugated (short spin), and placed at 4°C overnight. The total tau detection antibody (Millipore-Sigma, clone: 2A1-2E, mouse) was diluted 1:1000 in 1% Blocker A + 0.05% Tween 20. Goat anti-mouse sulfo-tagged antibody (Mesoscale Discovery) was diluted 1:500 in 1% Blocker A + 0.05% Tween 20. Coated assay plates were washed three times with 1X PBS + 0.05% Tween 20. Then, 150 µL/well of 5% Blocker A (Mesoscale Discovery R93AA-1) was added, plates were sealed and placed on an orbital shaker at room temperature for 2h. After blocking, plates were washed three times, followed by careful transfer of 100 µL/well of cell lysate to MSD plate. MSD plates were sealed again and placed on the orbital shaker at 4°C overnight. Next day, plates were washed with 1X PBS + 0.05% Tween 20 and incubated with 50 µL/well of detection antibody while shaking at room temperature for 2h. Next, plates were washed three times and incubated with 50 µL/well of anti-mouse SULFO-tagged antibody (Mesoscale Discovery) while shaking at room temperature for 1h. Plates were washed three times, and 150 µL/well of 1X MSD read buffer was added. Plates were immediately read on the MSD S600 instrument.

### Human iPSC-derived NPC culture and differentiation

Work with human induced pluripotent stem cell (iPSC) and derived neural progenitor cell (NPC) lines was performed under the Massachusetts General Hospital/Mass General Brigham-approved IRB Protocols 2010P001611 and 2022P001361. Reprogramming of patient dermal fibroblasts into iPSCs by non-integrating methods and subsequent conversion into cortical-enriched neural progenitor cells (NPCs) were previously described (*15, 60, 64*). The patient cell lines employed in this study were as follows: NPC line MGH2046-RC1 (*35, 75*), derived from a female individual with FTD carrying the autosomal dominant mutation tau-P301L (rs63751273); NPC line 300.12 (*15, 60*) was obtained from NSCI (peripheral blood mononuclear cells deposited in NCRD: https://ncrad.iu.edu/) and derived from a female individual with unknown diagnosis, carrying the splicing missense mutation tau-S305N (rs63751165); and NPC line MGH2069-RC1 (*35, 75*), which was derived from an unaffected-healthy (carrying wild type tau) female individual.

NPCs were cultured in 6-well (Fisher Scientific Corning) or 96-well flat bottom, black (Fisher Scientific Corning 3904) plates, coated with poly-ornithine (20 μg/mL in water, Sigma P3655-50MG) and laminin (10 μg/mL in PBS, Sigma L2020-1MG), referred to as POL-coated plates. Cells were cultured in proliferative conditions with *Neural Progenitor Medium* (NPM, Stem Cell Technology 05833-kit) supplemented with 1X penicillin-streptomycin (Pen/Strep, Gibco 15140), and passaged with Accutase (Sigma A6964). For NPC differentiation (up to eight weeks), cells were first plated at an average density of ∼120,000 cells/cm^2^ in POL-coated plates with NPM medium supplemented with 1X Pen/Strep and 10 μM DAPT inhibitor (Tocris 2634) for 24h. DAPT is a γ-secretase and Notch signaling inhibitor and is used in an initial selection phase to promote neuronal differentiation from NPCs. Cell medium was then replaced with *Neuronal Differentiation Medium* (NDM) [Neurobasal medium (Life Technologies 21103-049), 1X B27- Plus (Thermo A3582801), 20 ng/mL BDNF (Stem Cell Technology 78005.1), 20 ng/mL GDNF (Stem Cell Technology 78058.1), 0.5 mM dibutyryl-cAMP (Tocris 1141), 1% Glutamax (Gibco 35050-062) and 1% Pen/Strep (Gibco 15140)], also supplemented with the selection agent 10 μM DAPT (Tocris 2634). Medium was replaced daily for 7 days. At this point, and to promote further neuronal differentiation and maturation, cell medium was switched to *Complete BrainPhys medium* [BrainPhys Neuronal Medium (Stem Cell Tech. 005790), 1X B27-Plus (Thermo A3582801), 1X N2-Supplement-A (Stem Cell Technology 07152), 0.02 μg/mL BDNF (Stem Cell Technology 78005.1), 0.02 μg/mL GDNF (Stem Cell Technology 78058.1), 1 mM dibutyryl-cAMP (Tocris 1141), 200 nM ascorbic acid (Stem Cell Technology 72132), 1X Pen/Strep (Gibco 15140)]. Neuronal cultures were “fed” every 2.5 days with half medium/well volume changes, until time of treatment and/or collection. For generating neuronal cultures, we utilized tau-P301L NPCs (FACS purified CD133^+^/CD184^+^/CD271^−^) at passage 30–35; and tau-S305N NPCs (non- FACSed) at passage 4. Differentiated neurons in 6-well plates were used for mRNA and protein analysis. Differentiated neurons in 96-well plates were used for viability and stress rescue assays.

### Compound treatment of human iPSC-derived neurons

Compound treatment of neuronal cultures in 6-well plates was performed in 2 mL medium volume (*Complete BrainPhys*) by removing 1 mL of cell medium and adding 1 mL of new medium pre- mixed with the compound at the appropriate 2X concentration, followed by incubation at 37°C for the designated time. Compound treatment of neuronal cultures in 96-well plates was performed in 100 μL medium by adding compound directly to each well, followed by incubation at 37°C. Stock concentrations in serial dilutions were prepared fresh before each experiment, so that the volume of compound in DMSO added per well remained constant (≤ 0.1% DMSO). We utilized two treatment regimens: a 96h treatment during week 8 of differentiation, with the compound added to medium at the time of culture feeding; and a weekly treatment from weeks 5 to 8 of differentiation, i.e., a 3-week regimen with the compound added to medium once per week during culture feeding (with the second feeding of the week being done as usual, with BrainPhys medium).

### Neuronal viability assay

NPCs were plated at an average density of 150,000 cells/cm^2^ in black 96-well plates with a clear flat bottom (Fisher Scientific Corning 3904), differentiated for 5–7 weeks, and treated with compound for 3 weeks or 96h (Fig. 2A). Viability was measured at the end of week 8 with the alamarBlue HS cell viability assay (ThermoFisher Scientific A50101). AlamarBlue reagent was added at a 1:10 dilution in 100 μL cell medium, and incubated for 4h at 37°C. Readings were done in the Envision Multilabel Plate Reader (Perkin Elmer), according to the manufacturer’s instructions. Cell viability calculations were performed in Microsoft Excel (v.16.94) and graphs were plotted in GraphPad Prism 10.

### RT-qPCR analysis of neuronal 4R MAPT

Neurons were washed in DPBS (Corning), lifted into suspension by scraping, transferred to eppendorf tubes, and centrifugated at 3,000 *xg* for 5 min. The cell pellet was stored at -80°C until sample processing. Isolation of RNA and cDNA synthesis from iPSC-neuronal cultures was performed as described above for HEK293T cells. The mRNA levels of *4R MAPT* and *GAPDH* were quantified by RT-qPCR analysis using the same methodology described for the HEK293T cells. To amplify the *4R MAPT* we utilized : Taq_4R_qPCR forward 5’-GCGGGAAGGTGCAGATAATT-3’; Taq_4R_qPCR reverse 5’-GCTCAGGTCAACTGGTTTGT-3’; Taq_4R_qPCR probe 5’-AAGAAGCTGGATCTTAGCAACGTCCAGTCCA-3’. To amplify *GAPDH* we utilized Taq_hGADPH_qPCR forward 5’-CAACGGATTTGGTCGTATTGG-3’; Taq_hGADPH_qPCR reverse 5’-TGATGGCAACAATATCCACTTTACC-3’; Taq_hGADPH_qPCR probe 5’- CGCCTGGTCACCAGGGCTGCT-3’. Fold changes were calculated relative to vehicle (DMSO) treated sample and normalized to *GAPDH*. Values were plotted in GraphPad Prism 10.

### iPSC-derived neurons protein extraction and western blot analysis

Neurons were washed in DPBS (Corning), lifted into suspension by scraping, transferred to eppendorf tubes, and centrifugated at 3,000 *xg* for 5 min. Cell pellets were lysed in RIPA buffer (Boston Bio-Products BP-115) supplemented with 2% SDS (Biorad 1610418), 1% Halt Protease/Phosphatase inhibitors (Thermo Fisher Scientific 78442), 1:5000 Benzonase (Sigma E1014), and 10 mM DTT (NEB 7016L), for 15 min at room temperature. Lysates were centrifugated at 20,000 ×*g* for 20 min and the supernatants were transferred to new eppendorf tubes for analysis. Protein concentration quantification was performed using the Pierce BCA Protein Assay Kit - Reducing Agent Compatible (ThermoFisher Scientific 23252). For western blot, 10–20 μg of total protein per well, diluted in SDS blue loading dye (NEB B7703S) and heated to 95°C for 5 min were analyzed by SDS-PAGE. Electrophoresis was performed with the Novex NuPAGE Gel System (Invitrogen). Proteins were transferred from the gel onto PVDF membranes (Immobilon-P Membrane 0.45 µm–Millipore/Sigma IPVH00010) using standard procedures. Membranes were blocked in 5% BSA (Sigma A7906) in Tris-buffered saline with Tween-20 (TBST, Boston Bio-Products IBB-181), incubated overnight with primary antibody in 5% BSA- TBST at 4°C, followed by incubation with the corresponding HRP-linked secondary antibody at 1:4000 dilution (Cell Signaling Technology). Blots were developed with SuperSignal West Pico or Dura Chemiluminescent Substrate (Thermo Fisher Scientific 34580/34076) according to manufacturer’s instructions, exposed to autoradiographic films (LabScientific by ThermoFischer), and scanned on an Epson Perfection V600 Photo Scanner. Protein bands’ densitometry was measured with ImageJ (v. 1.53t) and normalized to the respective internal control (β-Actin) band. Calculations were performed in Microsoft Excel (v.16.94), and graphs were plotted in GraphPad Prism 10. The antibodies used were as follows: 4R tau (Abcam ab218314, Cell Signaling Tech. 79327s, Millipore 05-804), 3R tau (Millipore 05-803), total tau TAU5 (Invitrogen AHB0042), pTau-Ser396 (Invitrogen 44752G), pTau-Ser202/Thr205 (AT8, Thermo Scientific MN1020), pTau-T181 (AT270, Thermo Fisher Scientific MN1050), pTau-T231 (AT180, Thermo Fisher Scientific MN1040), SYN1 (Cedarlane Labs 106011(SY)), PSD95 (Cedarlane Labs 124011 (SY)), NeuN (Thermo Fisher Scientific 702022), β-III-Tubulin (TUJ1, Sigma T-8660), β-ACTIN (Sigma A1978). All the employed tau antibodies have been extensively characterized by us and others. For 4R tau detection we utilized the averaged densitometry across the three antibodies tested and show the individual results when appropriate in supplementary information.

### iPSC-derived neurons ICC and imaging

Neurons were differentiated as described above, at a starting cell density of 90,000 cells/cm^2^ in black, clear flat bottom, POL-coated 96-well plates (Corning) and differentiated for 7 weeks, followed by 96h compound treatment. Neurons were fixed with 4% (v/v) formaldehyde-PBS (Tousimis) for 30 min, washed in PBS (Corning), and incubated in blocking/permeabilization buffer [10 mg/mL BSA (Sigma), 0.05% (v/v) Tween-20 (Bio-Rad), 2% (v/v) goat serum (Life Technologies), 0.1% Triton X-100 (Bio-Rad), in PBS] for 2h. Neurons were incubated with primary antibodies overnight: PHF1 (kindly gifted by Dr. Peter Davies, Albert Einstein College of Medicine, NY USA) at 1:1000, MAP2 (Millipore/Chemicon AB5543) at 1:1000, Hoechst-33342 (Thermo Fisher) at 1:2500. Cells were washed with PBS and incubated with the corresponding AlexaFluor-conjugated secondary antibodies at 1:500 dilution (Life Technologies). Image acquisition was done with the INCell 6000 (GE Healthcare) automated confocal microscope.

### Protein solubility assay

For the protein solubility assays, cell lysis and protein fractionation based on detergent solubility were performed as previously described (*64, 69, 70*). Briefly, higher solubility proteins (S fractions) were purified in 1% Triton buffer [1% Triton X-100 (Thermo Fisher Scientific), 1% Halt Protease/Phosphatase inhibitors (Thermo Fisher Scientific 78442), 1:5000 Benzonase (Sigma E1014) and 10 mM DTT (NEB 7016L) in DPBS], whereas lower solubility pelleted proteins (P fractions) were solubilized in 5% SDS buffer [5% SDS (Biorad 1610418), 1% Halt Protease/Phosphatase inhibitors, 1:5000 Benzonase and 10 mM DTT in RIPA buffer (Boston Bio- Products BP-115)]. SDS-PAGE western blot was performed by loading 20 μg of each S-fraction and equal volume of the P-fraction onto pre-cast Tris-Acetate SDS-PAGE (Novex, Invitrogen). Western blot was performed as before. Protein bands’ densitometry values were measured with ImageJ (v. 1.53t) and normalized to the respective GAPDH intensity in the S-fraction. Calculations were done in Microsoft Excel (v.16.94) and graphs were plotted in GraphPad Prism 10.

### Neuronal stress vulnerability assay

To measure neuronal vulnerability, i.e., changes to cell viability in the presence of the stressor glutamate, we executed assays as previously described (Fig. 4F) (*64*). NPCs were plated (∼150,000 cells/cm^2^) and differentiated in 96-well plate format, for seven weeks. PTC compounds or DMSO (Vehicle, Sigma D2438) were added directly onto the media and incubated for 96h at 37°C (compound pre-treatment) in 200 μL/well of medium. Then, each well was treated with glutamate (Sigma G1251) at the specified final concentrations or DMSO (vehicle), by adding the solution directly to the medium, and incubated at 37°C for 18h (stressor treatment). At this time, viability was measured with the alamarBlue HS Cell Viability assay (ThermoFisher Scientific A50101). AlamarBlue reagent was added at a 1:10 dilution in 200 μL cell medium, and incubated for 4h at 37°C. Readings were done in the Envision Multilabel Plate Reader (Perkin Elmer), according to the manufacturer’s instructions. Cell viability calculations were performed in Microsoft Excel (v.16.94) and graphs were plotted in GraphPad Prism 10.

### Tauopathy mouse model and compound treatment

All animal studies were conducted at an *Association for Assessment and Accreditation of Laboratory Animal Care (AAALAC)*-approved facility, under an institutional animal care and use committee (IACUC)-approved protocol, in accordance with NIH guidelines. Study designs utilized *Animal Research: Reporting of In Vivo Experiments (ARIVE)* guidelines (https://arriveguidelines.org/arrive-guidelines). The mice were housed in the animal facility at Robert Wood Johnson Medical School, provided with access to food and water ad libitum, and maintained on a 12-hour light/dark cycle. Animals in which the endogenous mouse *Mapt* gene is replaced by the human MAPT gene carrying the N279K mutation (gene-replacement/GR-mouse model – *MAPT*(H1.0*N279K)-GR (JAX #35794)) were obtained from Dr. Michael Koob (University of Minnesota) (*61*). For our studies we employed wildtype (*MAPT*-GR WT(H1)), heterozygous tau-N279K/WT, and homozygous tau-N279K/N279K animals. Two studies were performed. In the first study, 2-month-old N279K mice were dosed via oral gavage with 7 mg/kg PTC-003 (at 10 mL/kg in 0.5% HPMC with 0.1% Tween 80) in combination with 50 mg/kg aminobenzotriazole (ABT), a pan inhibitor of the cytochrome P450 (*71*). At 1h, 2h, 4h, 7h, 16h, and 24h after dosing, mice were euthanized and plasma and brain tissue were collected for subsequent analysis of drug levels and *3R/4R MAPT* mRNA levels. In the second study, 2-month-old homozygous tau-N279K/N279K mice were dosed daily via oral gavage with ABT (50 mg/kg) 2h prior to dosing with vehicle or PTC-003 (1 or 2.5 mg/kg at 10 mL/kg in 0.5% HPMC with 0.1% Tween 80) for 21 days. At necropsy, the brains were collected for analysis of 3R tau, total tau and pTau231 protein levels.

### RNA isolation and qRT-PCR analysis of MAPT transcripts in mouse tissue

Mouse brains were snap-frozen in liquid nitrogen upon collection. Tissues (full brain) were homogenized in QIAzol Lysis Reagent (Qiagen 79306) using a TissueLyser II (Qiagen). Total RNA was extracted using the RNeasy Lipid Tissue kit (Qiagen 74804) following the protocol provided by the manufacturer. The yield, purity, and quality of the total RNA for each sample were determined using a Nanodrop ND-8000 spectrophotometer. The mRNA levels of *3R MAPT*, *4R MAPT* and *GAPDH* were quantified by qRT-PCR analysis using CFX384 Real-Time System (BioRad) and Taqman-based probes (ThermoFisher Scientific) following the same protocol described above for HEK293T and neuronal cells.

### MSD immunoassay for 3R tau and pTau231 protein quantification in mouse tissue

Tissue samples were collected, snap-frozen in liquid nitrogen, weighed and homogenized on the TissueLyzer II (Qiagen) in MSD tris lysis buffer (MesoScale Discovery, R60TX), supplemented with 1X Halt Protease inhibitor (ThermoFisher, 78438), 1X Phosphatase Inhibitors II and III (Sigma-Aldrich, P5726 and P0044), 1 mM PMSF (Sigma-Aldrich, 93482) and 10 mM NaF (Sigma-Aldrich, BP-450) at a tissue-weight to lysis buffer volume of 100 mg/mL. The samples were then centrifuged at 4°C for 20 min at 14,000 *xg* in a microcentrifuge. The homogenates were transferred into microcentrifuge tubes and stored at -80**°**C. Total protein concentration was determined using the Pierce BCA Protein Assay Kit (ThermoFisher, 23225). Samples were run in duplicate and averaged. 3R tau was quantified using the same MSD protocol described for quantification in cells. For measuring pTau231 in tissue homogenates, we used MSD multi-spot Phospho (Thr231)/Total tau assay kit (MesoScale Discovery, K15121D-2) and followed manufacturer’s provided protocol.

### Statistical Analysis

Graphed data represent mean values ± SD (standard deviation) or ± SEM (standard error of the mean), calculated using Microsoft Excel and GraphPad Prism. *P*-value < 0.05 was considered the threshold for statistical significance. *P*-value significance intervals (*) are provided within each figure legend, together with the statistical test performed for each experiment. *N* values are indicated within figure legends and refer to biological replicates (NPC differentiation and neuronal cultures independently setup and analyzed at different times or number of individual animals treated/tested), whereas technical replicates refer to repeated analysis of the same samples. Derived statistics correspond to analysis of averaged values across biological replicates.

## Acknowledgments

We wish to thank members of the Tau Consortium (Rainwater Charitable Foundation) for helpful feedback on tauopathy iPSC-derived cell models and *MAPT* splicing iPSC-neuronal models. The Tau-S305N NPC line was acquired from The Neural Stem Cell Institute (NSCI). We thank the laboratory of Dr. Peter Davies (Albert Einstein College of Medicine, NY USA) for kindly sharing the tau PHF1 antibody.

## Author contributions

Conceptualization and design: MCS, JT, CRT, MW, EM

Methodology and analysis: MCS, HL, PP, JL, YY, CM, MA, SJB, KS, MD, AM, NZ, JN, CRT, MW, EW

Project supervision, data interpretation: MCS, JT, EM

Funding acquisition: MCS, EM

Resources: SL, TB, ST, SJH, MK PTC program coordinator: KD

Writing – original draft: MCS, EM

Writing – review and editing: MCS, HL, PP, JL, SJB, MD, JN, TB, ST, SJH, MK, MW, EM

## Data and materials availability

The main data supporting the findings of this study are available within the article and its Supplementary Figures. Additional raw data and protocols that support the findings within the article are available from the corresponding authors upon reasonable request.

## LIST OF SUPPLEMENTARY MATERIALS

Supplementary Figures – Fig. S1 to Fig. S7.

