## Supplementary Figures for "*MAPT* Splicing Modulators as a Therapeutic Strategy for Tauopathies"

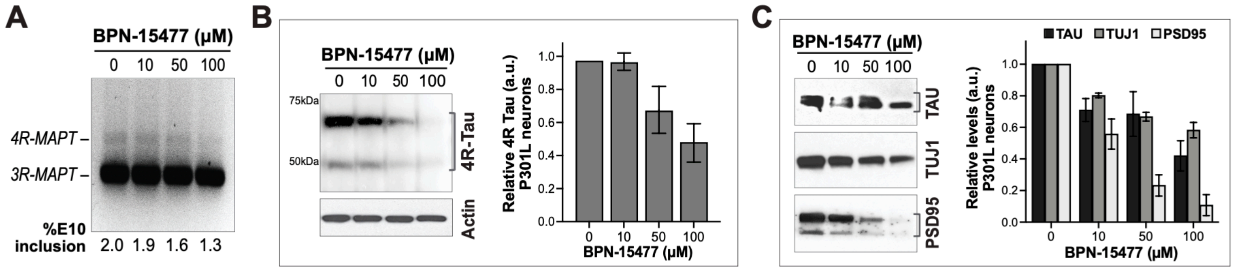


**Figure S1. BPN-15477 promotes *MAPT* exon 10 exclusion in tau-P301L neurons (related to Fig. 1).** Dose-dependent effect of BPN-15477 on endogenous *MAPT* splicing in human tau-P301L neurons treated from week 3 to week 8 of differentiation. BPN-15477 promotes exon 10 skipping (–E10), resulting in **(A)** reduced *4R MAPT* expression, as shown by RT-PCR analysis, and **(B)** decreased 4R tau protein levels, as shown by western blot analysis. **(C)** Neuronal morphology (β-III-tubulin/TUJ1) and synaptic (PSD95) markers show dose-dependent levels' reduction, in parallel to the decrease in total tau (TAU5 antibody). This reduction likely reflects a loss in neuronal integrity, and potential compound toxicity. Representative gels/blots are shown, and bar graphs represent mean densitometry values (normalized to actin) relative to vehicle DMSO-treated neurons (0 μM) ± SD; $N = 2$.

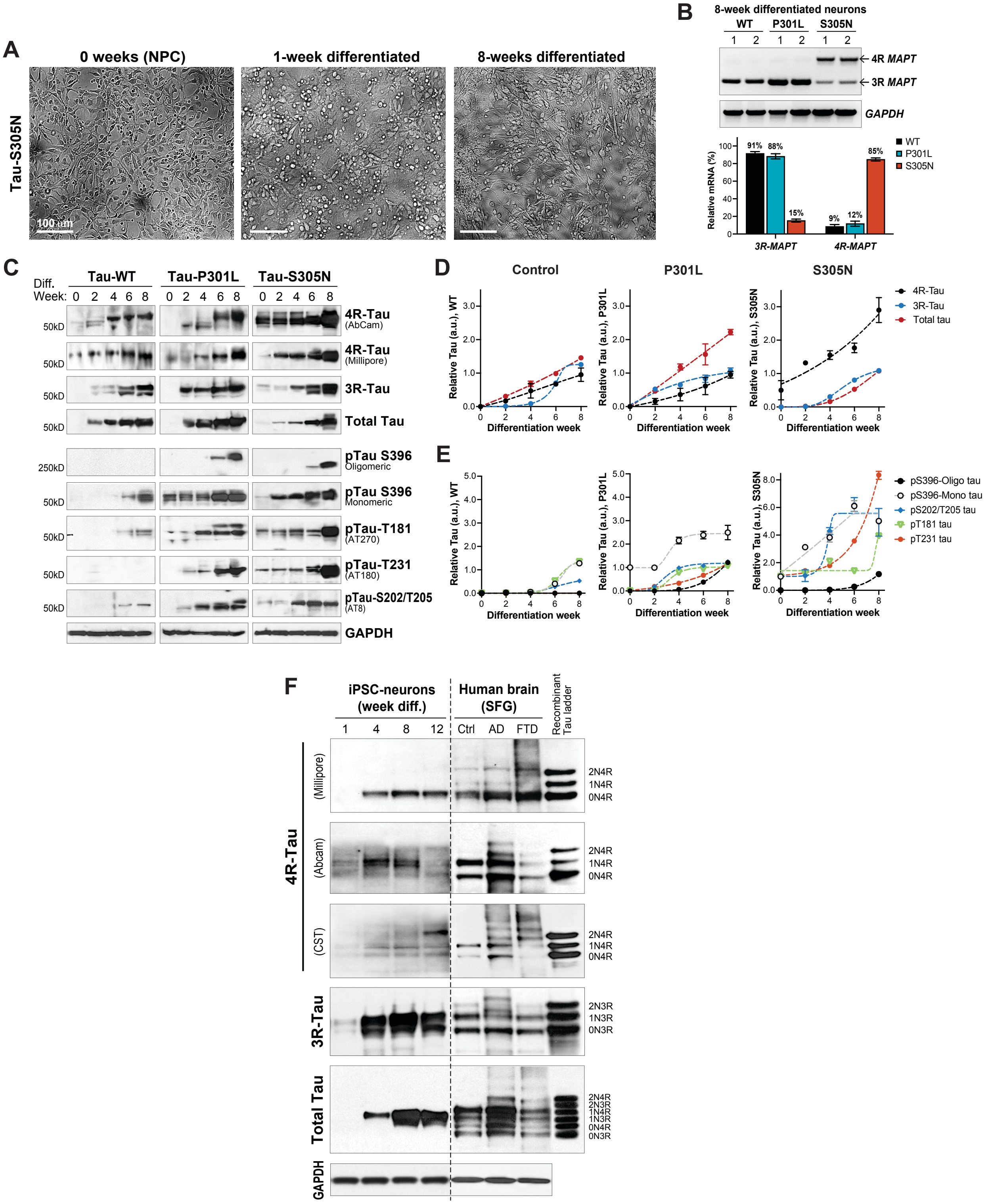


**Figure S2. Characterization of tauopathy phenotypes in the tau-S305N iPSC-derived neuronal model and antibody validation in human brain tissue (related to Fig. 2). (A)** The tau-S305N cell line (Id 300.12) was differentiated from NPC stage with BrainPhys medium. Bright-field microscopy images capture the progression in neuronal processes maturation and network formation within the first 8 weeks of differentiation. Scale bar 100 µm. **(B)** Representative RT-PCR analysis of *MAPT* splicing in P301L, S305N and control wild type (WT) neurons, demonstrating increased exon 10 inclusion and *4R MAPT* expression driven by the splicing mutation in S305N neurons. In P301L neurons the level of *4R MAPT* detection is low (~12% of total *MAPT*). Bar graphs represent mean densitometry (%), normalized to GAPDH ± SD (*N*

= 2). **(C-E)** Comparative analysis of tau protein markers across three genotypes (WT, P301L, S305N) during a time-course of neural differentiation between week 0 (NPC) and week 8, by western blot and densitometry analysis. The markers measured include 4R tau, 3R tau, total tau (TAU5 antibody), as well as phospho-tau (pTau) species associated with human brain pathology in FTD, including pTau-S396 (which detects monomeric ~50kDa tau and oligomeric >250 kDa tau), pTau-T181 (AT270 antibody), pTau-T231 (AT180 antibody) and pTau-S202/T205 (AT8 antibody). Time-dependent increase in tau is shown (D, E) by densitometry quantification, normalized to actin and relative to NPC stage ± SD ($N$ = 2 biological replicates). **(F)** Validation of tau antibodies. We utilized iPSC neurons at 1, 4, 8 and 12 weeks of differentiation (P301L) and protein lysates from brain tissue (superior frontal gyrus, SFG) from a control-healthy individual, an Alzheimer's disease (AD) patient and an FTD patient carrying the P301L mutation. Recombinant tau ladder (*rightmost lane*) was used for band MW guidance. We were able to validate three commercial 4R tau antibodies to use in our study and used 3R tau and total tau detection as controls for comparison between human iPSC neurons and brain tissue.

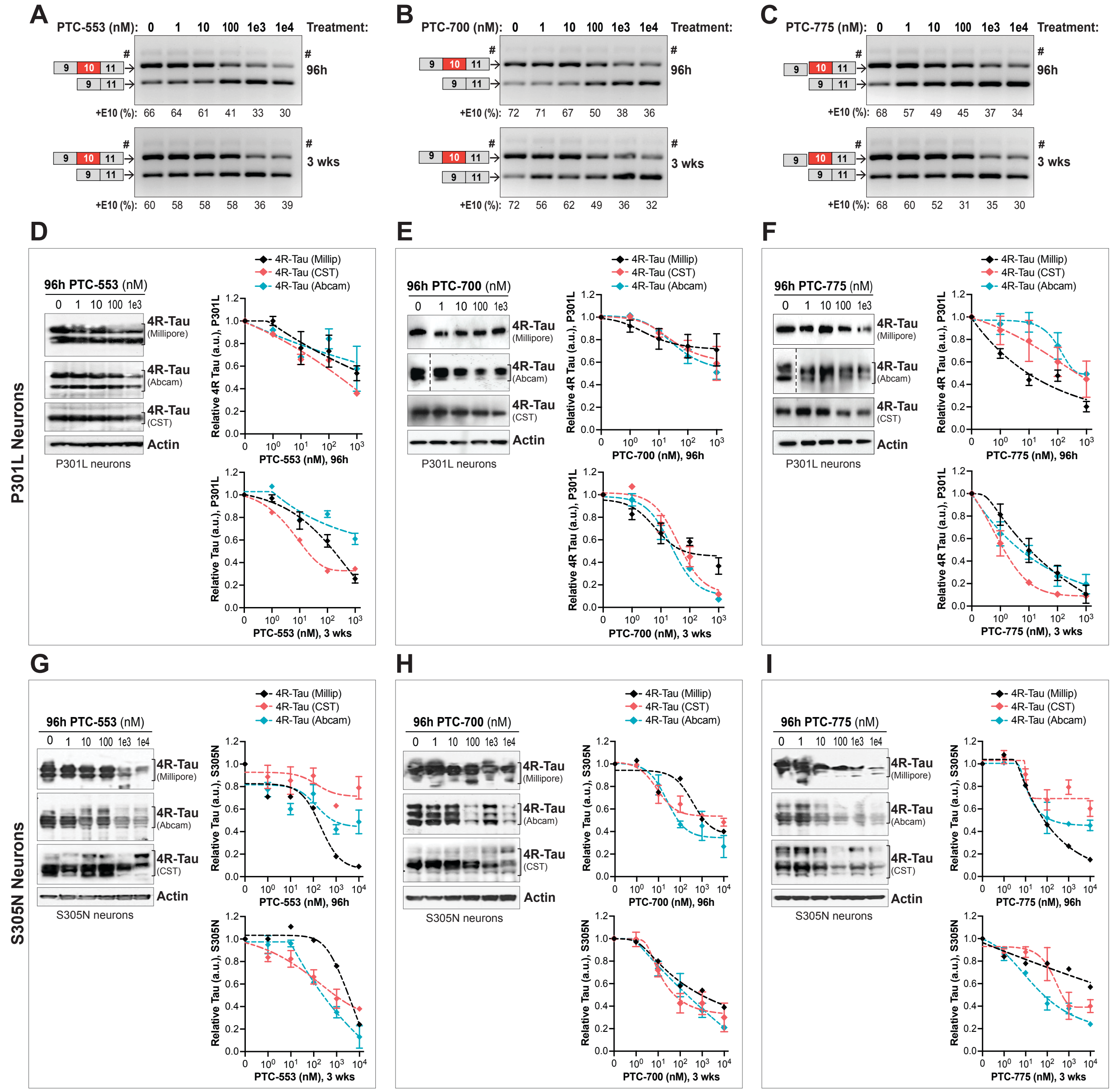


**Figure S3. SMCs promote *MAPT* exon 10 skipping and reduce 4R tau protein levels in P301L and S305N neuronal models (related to Fig. 2). (A-C)** Representative RT-PCR gel electrophoresis analysis of *MAPT* splicing in S305N neurons treated with PTC-553 (A), PTC-700 (B), and PTC-775 (C) for 96h or 3 weeks, showing a dose-dependent increase in *3R MAPT* (–E10, lower band) and a corresponding decrease in *4R MAPT* (+E10, upper band). A faint higher band (#) is also visible and shares the same nucleotide sequence as the *4R MAPT* band, likely representing a secondary structure of the PCR product. **(D-F)** Western blot analysis of SMC dose-curves in P301L neurons after 96h and 3 weeks of treatment, with individual 4R tau antibodies quantification (Millipore, Abcam, Cell Signaling Technology/CST). Representative blots for the 96h treatments are shown for PTC-553 (A), PTC-700 (B) and PTC-775 (C). Graph data points represent mean densitometry relative to vehicle (DMSO)-treated neurons ± SEM, with N ≥ 3 biological replicates. Dashed lines on blots (B) and (C) indicate cropped images solely for the purpose of this figure to exclude data unrelated to this study, all samples were run in the same gel. **(G-I)** Western blot analysis of SMCs dose-curves in S305N neurons, after 96h and 3 weeks of treatment, with representative blots shown for 96h compound treatment. Densitometry analysis of 4R tau levels, relative to vehicle-treated neurons (normalized to actin) is shown for each 4R antibody tested, with data points representing mean densitometry ± SEM, with N ≥ 3 biological replicates.

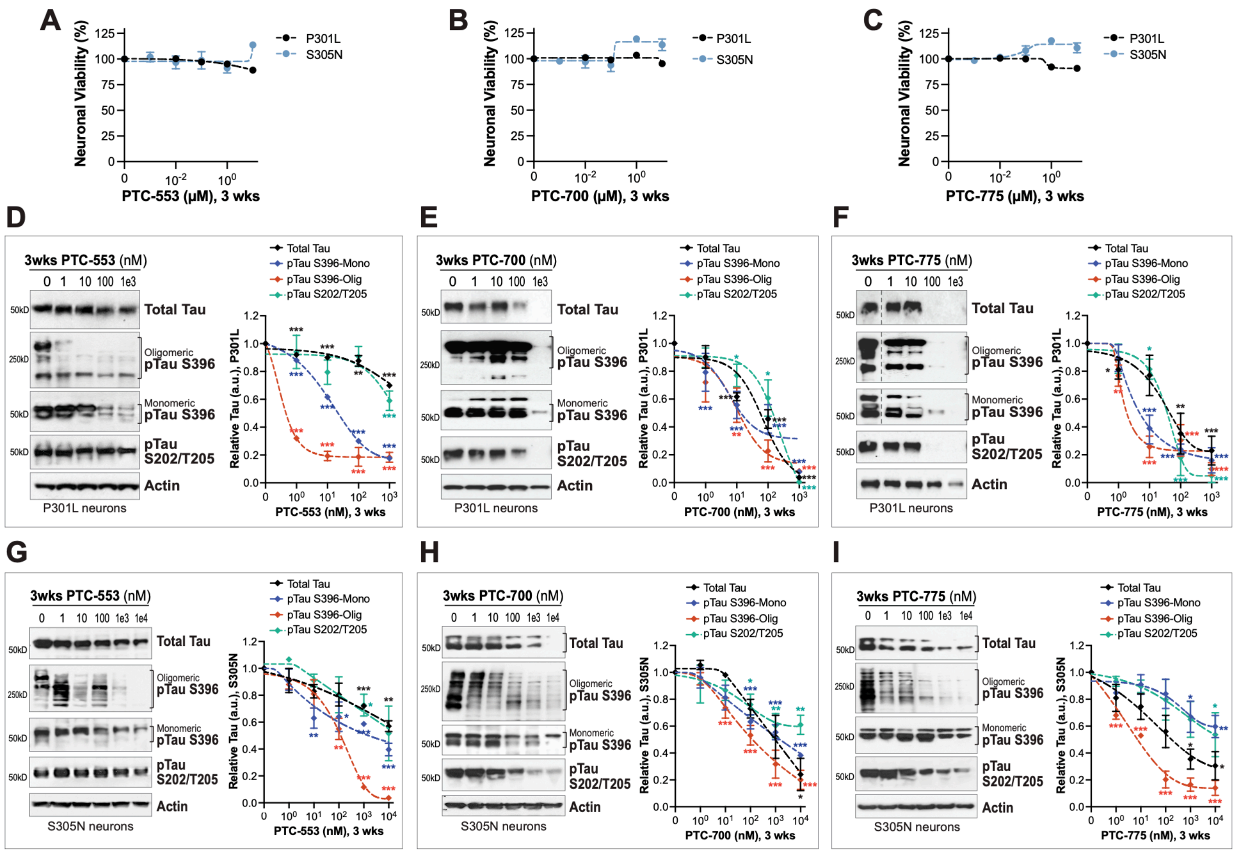

**Figure S4. SMC treatment of tauopathy neurons for 3 weeks reduces accumulation of total tau and pTau without compromising neuronal viability (related to Fig. 3). (A-C)** Viability dose curves for 8-week differentiated P301L and S305N neurons treated with PTC-553 (A), PTC-700 (B) or PTC-775 (C) for 3 weeks. Data points represent mean viability relative to vehicle-treated neurons (100% viability) ± SEM, with $N = 3$ biological replicates. **(D-F)** SMCs dose curves for total tau (TAU5 antibody), pTau-S396 (high MW–oligomeric >250 kDa and monomeric ~50kDa tau), and pTau-S202/T205 (AT8 antibody) after 3 weeks of treatment of P301L neurons, by western blot and densitometry analysis. Dashed lines in blots (F) indicate cropped images solely for the purpose of this figure to exclude data unrelated to this study, all samples were run in the same gel. **(G-I)** Corresponding analysis for S305N neurons. Representative blots are shown for 3-week treated neurons. Data points represent mean densitometry relative to vehicle-treated neurons ± SEM, with $N = 3$ biological replicates. Statistical analysis was performed using a two-tailed, paired Student's t-test for each compound concentration vs. vehicle (0 μM). Significance thresholds: $*p \leq 0.05$, $**p \leq 0.01$, and $***p \leq 0.001$.

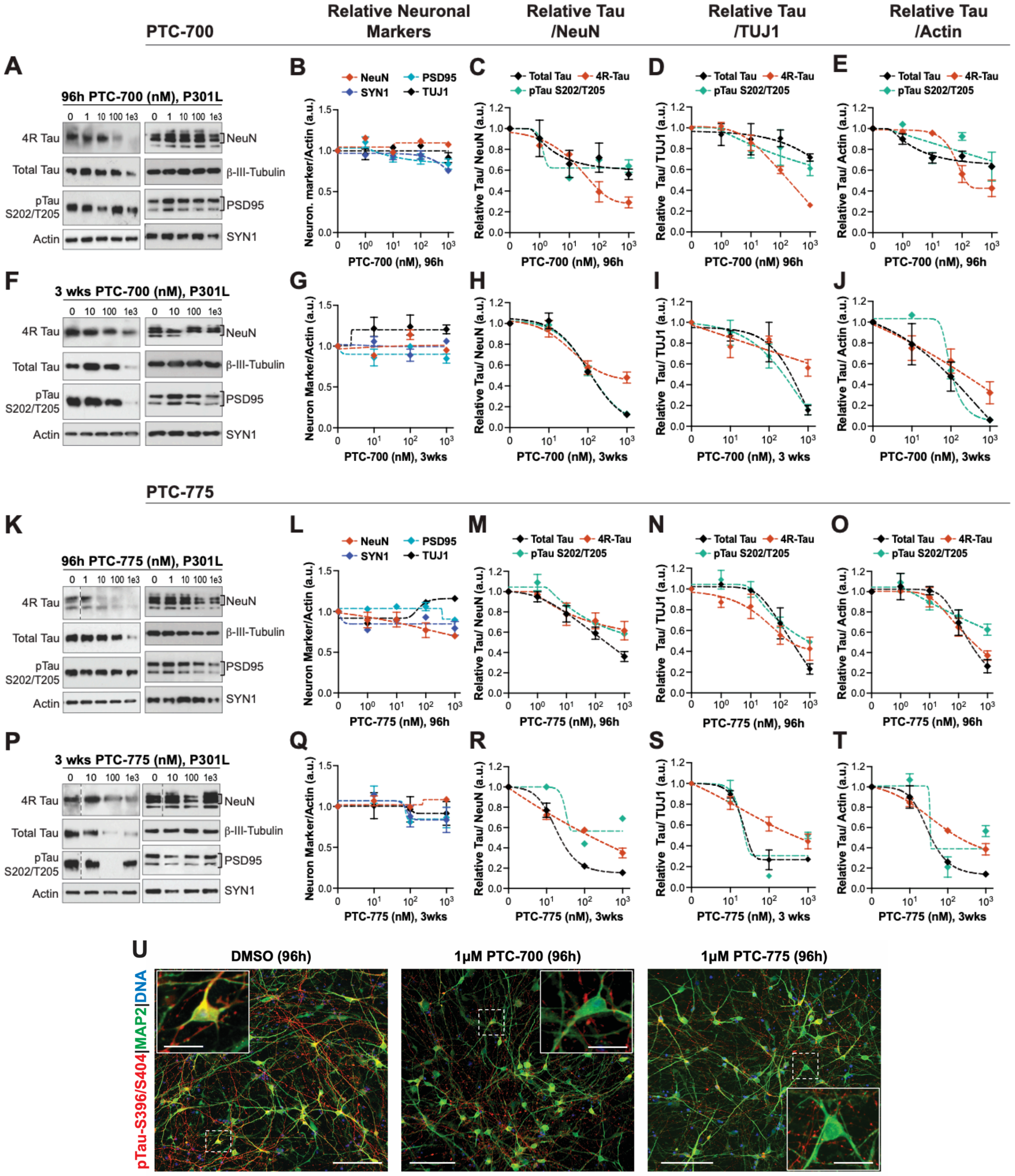


**Figure S5. Evaluation of neuronal integrity markers in P301L neurons treated with PTC-700 and PTC-775 (related to Fig. 3).** Tau-P301L neurons were treated with PTC-700 for 96h **(A-E)** or 3 weeks **(F-J)**, and with PTC-775 for 96h **(K-O)** or 3 weeks **(P-T)**. Neuronal lysates were analyzed by western blot and probed for neuronal markers (NeuN, β-III-tubulin/TUJ1, PSD95, SYN1) and tau markers (4R tau, total tau/TAU5 and pTau-S202/T205 (AT8)). **(B, G, L, Q)** Densitometry analysis of relative levels of neuronal markers normalized to actin and relative to vehicle/DMSO, showed that compound treatment did not affect neuronal integrity beyond SD variability, indicating no neuronal toxicity by the SMCs. This was also demonstrated by comparing changes in tau markers when normalized to NeuN **(C, H, M, R)** or normalized to TUJ1 (**D, I, N, S**) vs. normalized to actin **(E, J, O, T)**, which resulted in similar dose curves across normalization markers, supporting that SMCs did not affect number of neurons or neuronal integrity/morphology. The data points indicate mean relative densitometry ± SD, with $N = 2$ biological replicates (and 2 technical replicates). (**U**) Immunocytochemistry (ICC) of P301L neurons at 8 weeks of differentiation treated with vehicle (DMSO), PTC-700 or PTC-775 for 96h. Neurons were stained for pTau-S396/S404 in red (PHF1 antibody), MAP2 dendritic marker in green, and nuclei (hoechst-33342) in blue. Representative microscopy images (scale bar 100 μm) show the reduction in tau staining with SMC treatment, particularly obvious in the cell bodies (see insets, scale bar 25 μm), and demonstrate unchanged neuronal culture integrity upon treatment.

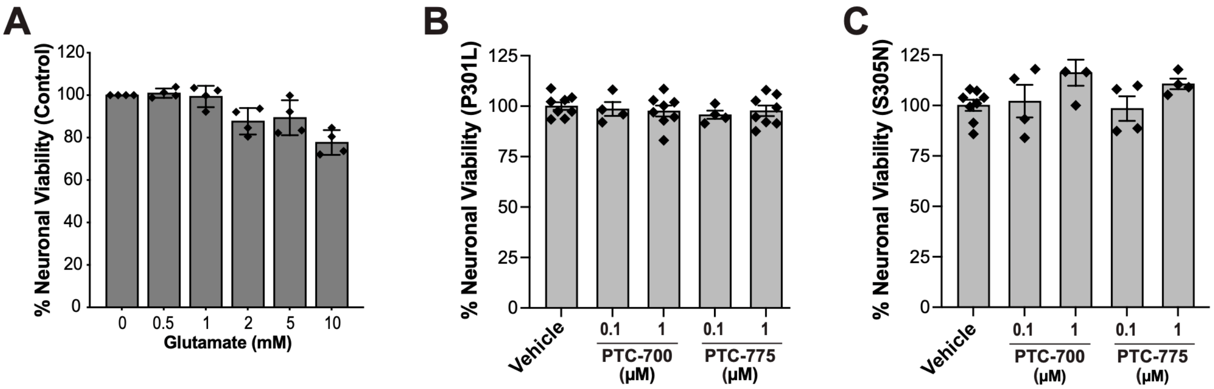


**Figure S6. Neuronal viability of tau-P301L and tau-S305N models under different compound treatment conditions (related to Fig. 4). (A)** Effect of increasing glutamate concentrations on the neuronal viability of tau-WT control neurons treated for 18h. Individual data points are indicated and graph bars represent mean % neuronal viability relative to vehicle (DMSO) treated neurons ± SEM (*N* = 4 biological replicates). **(B, C)** Neuronal viability for P301L (B) and S305N (C) neurons treated with SMC alone for 96h in the stress vulnerability assays. Individual data points are indicated and graph bars represent mean neuronal viability (%) relative to vehicle (DMSO) treated neurons ± SEM (*N* = 4 biological replicates with ± 2 technical replicates per assay).

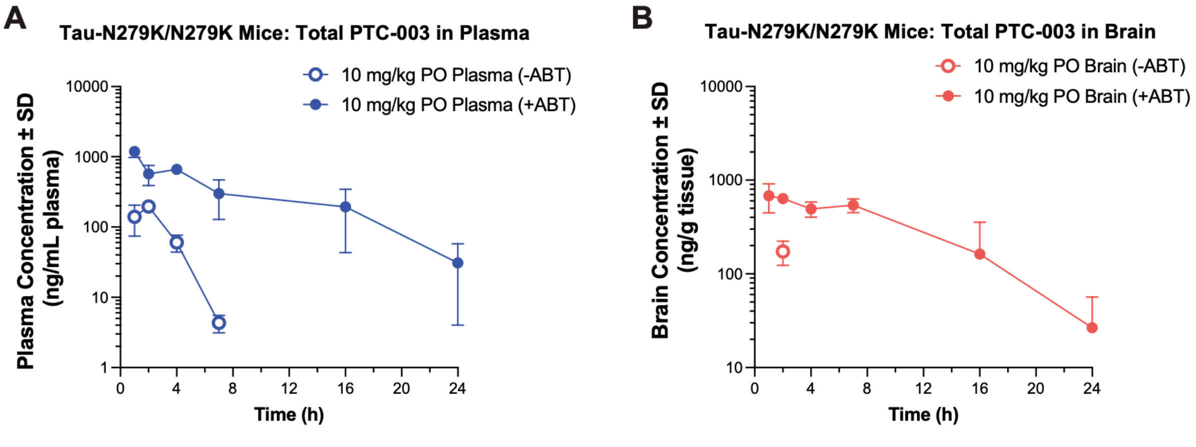


**Figure S7. Total PTC-003 levels in homozygous tau-N279K mice upon oral administration (related to Figure 6). (A)** Total plasma concentration of PTC-003 over a time course of 24h post-dosing, with (+) or without (–) ABT (1-aminobenzotriazole) co-administration. Compound levels were measured by mass spectrometry. **(B)** Brain tissue total concentration of PTC-003 over the same 24h time course, with (+) or without (–) ABT, measured by mass spectrometry.
